# Clonally expanded virus-specific CD8 T cells acquire diverse transcriptional phenotypes during acute, chronic, and latent infections

**DOI:** 10.1101/2021.06.29.450285

**Authors:** Raphael Kuhn, Ioana Sandu, Andreas Agrafiotis, Kai-Lin Hong, Daniel Neumeier, Doron Merkler, Annette Oxenius, Sai T. Reddy, Alexander Yermanos

## Abstract

CD8+ T cells play a crucial role in the control and resolution of viral infections and can adopt a wide range of phenotypes and effector functions depending on the inflammatory context and the duration and extent of antigen exposure. Similarly, viral infections can exert diverse selective pressures on populations of clonally related T cells. Technical limitations have nevertheless made it challenging to investigate the relationship between clonal selection and transcriptional phenotypes of virus-specific T cells. We therefore performed single-cell T cell receptor (TCR) repertoire and transcriptome sequencing of virus-specific CD8 T cells in murine models of acute, chronic and latent infection. We observed clear infection-specific populations corresponding to memory, effector, exhausted, and inflationary phenotypes. We further uncovered a mouse-specific and polyclonal T cell response, despite all T cells sharing specificity to a single viral epitope, which was accompanied by stereotypic TCR germline gene usage in all three infection types. Persistent antigen exposure during chronic and latent viral infections resulted in a higher proportion of clonally expanded T cells relative to acute infection. We furthermore observed a relationship between transcriptional heterogeneity and clonal expansion for all three infections, with highly expanded clones having distinct transcriptional phenotypes relative to lowly expanded clones. Finally, we developed and utilized a bioinformatic pipeline integrating pseudotime and clonality, termed Clonotyme, to further support a model in which expanded virus-specific CD8+ T cells adopt heterogenic, yet preferentially, effector-like phenotypes. Together our work relates clonal selection to gene expression in the context of viral infection and further provides a dataset and accompanying software for the immunological community.

## Introduction

T cells adopt a wide range of phenotypes and effector functions to orchestrate host-defense against infection. Viral infections can be loosely divided into acute and persistent (chronic and latent) infections, with influenza and SARS-CoV-2 being examples of the former and human immunodeficiency virus (HIV), cytomegalovirus (CMV), and hepatitis B virus examples of the later. Murine models of acute, chronic, and latent viral infections have been used to investigate the diverse phenotypes and functions of CD8+ T cells and have been instrumental in characterizing effector, memory, exhausted, inflationary, and self-renewing T cell populations (Frebel, Richter, and Oxenius 2010; Utzschneider et al. 2016; Wherry 2011; Sandu et al. 2020; Welten et al. 2020).

Laboratory studies with the lymphocytic choriomeningitis virus (LCMV) in mice have revealed that acute infections are characterized by the rapid recruitment and differentiation of virus-specific effector CD8+ T cells that enable viral clearance within days (Wherry et al. 2003). This is in contrast to chronic LCMV infections, where prolonged TCR stimulation results in the upregulation of inhibitory molecules and a decrease in effector capabilities, collectively termed T cell exhaustion (Wherry 2011). Finally, infection with another common mouse virus, murine cytomegalovirus (MCMV), has demonstrated to induce a population of expanded CD8+ T cells that respond to the latent reactivation events characteristic of herpes viruses, collectively termed inflationary T cells (Wiesel and Oxenius 2012; Karrer et al. 2003; Welten et al. 2020; Klenerman and Oxenius 2016). Reductionist approaches involving transgenic animals have been instrumental to characterize infection-specific T cell phenotypes, as transgenic CD8+ T cells expressing virus-specific TCRs can be transferred into naive hosts and profiled following viral infection (Sandu et al. 2020). While this approach is crucial to remove the possible variability between TCR affinities and avidities, it nevertheless introduces into the host an artificially high number of virus-specific CD8+ T cells expressing the same TCR. Similarly, as thousands of transferred TCR transgenic T cells are introduced into naive mice, it is challenging to relate clonal relationships to the dynamic phenotypes at the single-cell resolution.

While recent studies have leveraged bulk sequencing of the TCR beta (TRB) chain during acute, chronic, and latent murine infections (Welten et al. 2020; Yermanos, Sandu, et al. 2020; Chang et al. 2020), these methodologies are inherently limited by the inability to accurately access clonal expansion and further relate transcriptional profiles to those expanded TRB clones. Recent advances in single-cell immune repertoire sequencing can link the complete TCR beta and alpha sequence (VDJ) to gene expression (GEX) at the single-cell resolution (Yermanos, Neumeier, et al. 2021; Yermanos, Agrafiotis, et al. 2021; Horns, Dekker, and Quake 2020). This technology has recently demonstrated dynamic clonal and transcriptional profiles for virus-specific CD4+ T cells in the context of acute LCMV infection (Khatun et al. 2021), however, it remains unknown how previously described memory, effector, exhaustion, and inflationary phenotypes of virus-specific CD8+ T cells relate to antigen-driven clonal selection. We therefore performed single-cell TCR repertoire sequencing to investigate how the virus-specific CD8 T cell response varies across acute, chronic and persistent infections, which resulted in infection-specific transcriptional fingerprints. We additionally discovered a largely private and polyclonal T cell response in all three infection models, with chronic and latent infection showing higher levels of clonal expansion. Finally, our results indicate that clonally expanded T cells demonstrated transcriptional heterogeneity across all three infections, which was supported both by transcriptional clustering and pseudotime analysis.

## Results

### Single-cell sequencing recovers diverse transcriptional signatures of virus-specific CD8+ T cells

To profile the virus-specific CD8 T cell response, we leveraged three previously described models of murine viral infection, namely acute LCMV (low-dose clone 13), chronic LCMV (high-dose clone 13), and latent MCMV (clone MCMV-*ie2*-gp33) infection (Welten et al. 2015; Kräutler et al. 2020). An advantage of using these three viruses is that they all contain the gp_33-42_ (GP33) viral peptide epitope, which enables isolation of endogenous virus-specific CD8+ T cells using MHC-tetramers. While the GP33 peptide is naturally encoded in the LCMV genome, it has been engineered into the *ie2* gene locus of MCMV and gives rise to a population of virus-specific T cells termed inflationary T cells (Welten et al. 2020, 2015). We therefore isolated GP33-specific CD8+ T cells from the spleen of mice at 28 days post infection (dpi), separated into cohorts of either acute LCMV, chronic LCMV, or latent MCMV-*ie2*-gp33 infection. While we intended to include GP33-specific CD8+ T cells following peptide immunization with the GP33 peptide in CpG, we were unable to obtain sufficient numbers of GP33-specific cells 28 dpi and therefore excluded this group for future sequencing (Figure S1A). The virus-specific CD8+ T cells were then processed for single-cell sequencing of their TCR repertoires and transcriptomes by following the 10X genomics workflow (5’ immune profiling with V(D)J and GEX protocol) (Figures 1A, S1A). Following single-cell sequencing and alignment to the murine reference transcriptome, we recovered GEX information from thousands of virus-specific CD8+ T cells from each mouse (Figures 1B, 1C), with the median number of genes per cell ranging from 866 to 1210 for all mice (Figure 1D), accompanied by comparable percentages of mitochondrial genes and sequencing reads for all samples (Figures S1B, S1C).

**Figure 1.**
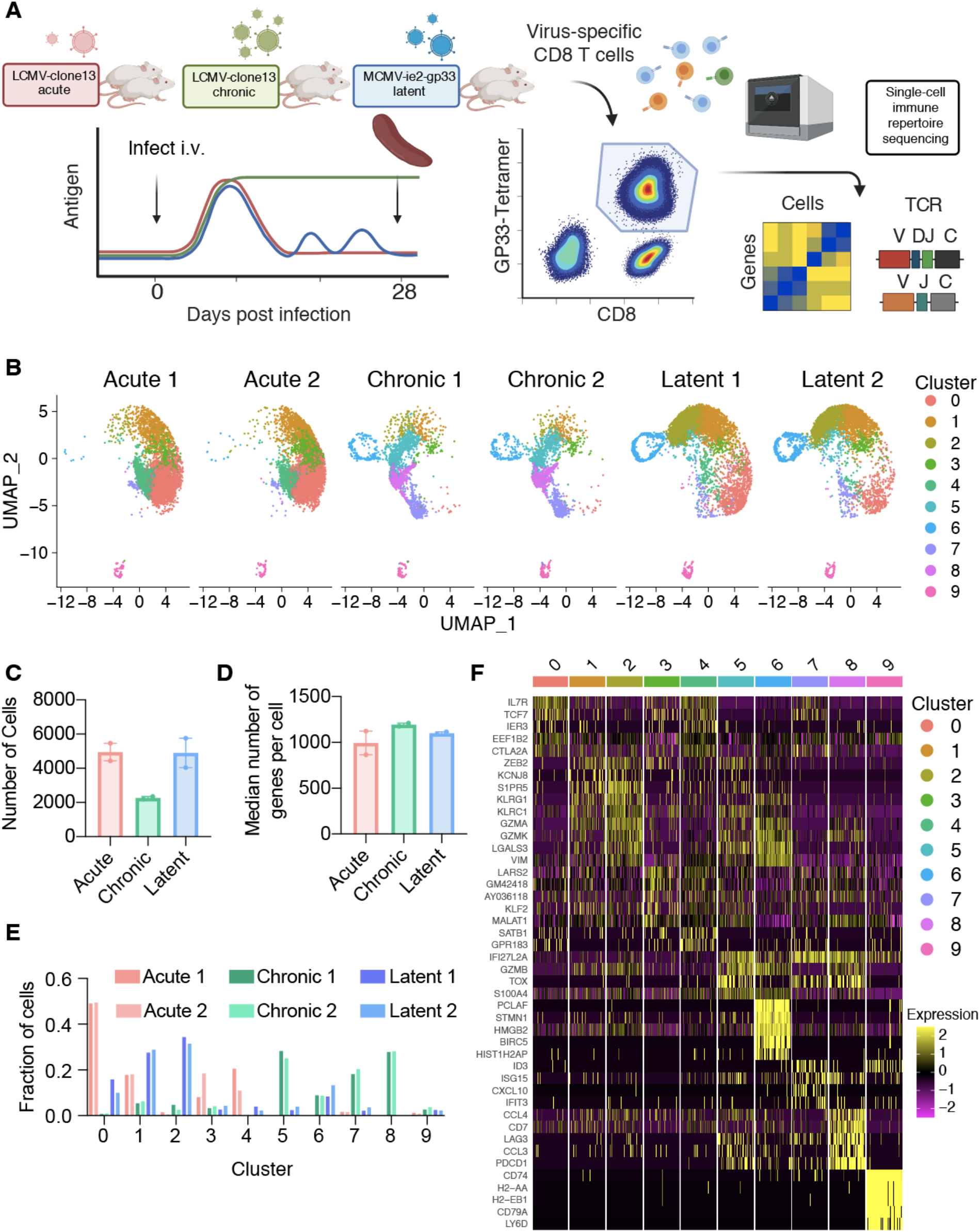
Single-cell immune repertoire sequencing recovers thousands of transcriptomes of virus-specific CD8+ T cells. A. Experimental overview. B. Uniform manifold approximation projection (UMAP) split by sample. Each point represents a cell and color corresponds to transcriptional clusters. All cells from all samples were integrated in this single UMAP. C. Number of cells with gene expression information for each mouse. D. Median number of genes per cell for each mouse. E. Fraction of cells in a particular transcriptional cluster for each sample. F. Top five significant genes defining each cluster ranked by average log fold change.

Acute, chronic, and latent infections have been reported to have distinct phenotypes of virus-specific CD8+ T cells, corresponding to memory, exhaustion, and inflationary subsets (Moskophidis et al. 1993; Wherry et al. 2007; Kaech et al. 2003; Kaech and Cui 2012; Klenerman and Oxenius 2016). Performing unsupervised clustering based on total gene expression [excluding genes relating to TCRs, i.e., V-, D-, J-, and constant region (C) genes] and subsequently visualizing the cells from all mice revealed infection-specific clustering (Figures 1B, S1D, S1E). Quantifying the proportion of cells in each cluster demonstrated that the transcriptional profiles were highly reproducible across biological replicates (Figures 1E, S1C), with CD8+ T cells found in distinct clusters for acute LCMV (clusters 0,1,3,4), chronic LCMV (5,6,7,8), and latent MCMV (0,1,2,6) infections. Further unbiased investigation into the most expressed genes per cluster revealed a plethora of genes previously reported in the context of viral infections, such as *Il7r*, *Tcf7*, *Zeb2*, *Klrg1*, *Gzma*, *Gzmb*, *Gzmk*, *Vim*, *Lgals3*, *Tox*, *Lag3*, *Pdc1*, *Id3*. Together, these patterns of gene expression suggested the presence of distinct memory (clusters 0, 3, 4), effector (clusters 1, 2, 5), exhausted (cluster 8), memory-like(cluster 7), proliferative (cluster 6) subsets, in addition to a small population of B cells present (cluster 9) in all samples, suggesting minor contamination (Figure 1F).

### Differential gene expression analysis, gene ontology, and gene set enrichment confirm memory, effector, exhaustion, and inflationary T cells

We next performed differential gene expression and calculated most up and down-regulated genes to determine if infection conditions would further separate transcriptional phenotypes (Figure 2A). Genes characteristic of T cell exhaustion were upregulated in the chronic LCMV infection (e.g., *Pdcd1*, *Tox*, *Lag3*), whereas genes associated with memory formation and inflationary phenotypes were upregulated in the acute LCMV (e.g., *Il7r*) and latent MCMV (e.g., *Klrg1*) infections (Figure 2A). As many of these genes have been previously described in the context of viral infection, we investigated whether the expression of additional genes commonly used to differentiate populations of CD8+ T cells could further differentiate infection types. Genes such as *Cd8a* and *Cd3e* were expressed ubiquitously across all cells (Figure S2), whereas exhaustion markers such as *Pdcd1*, *Tim3*, *Lag3*, and *Ctla4* were preferentially localized to the cells arising from chronic infection (Figures 2B, S2, S3). We additionally observed a population of cells from chronically infected mice coexpressing *Pdcd1* and *Tcf7* (Figures 2B, S2, S3), which has been described previously as “stem-like”, “memory-like” or “progenitor-exhausted”, and serves to sustain the effector and exhausted population in the context of chronic infection (Siddiqui et al. 2019; Utzschneider et al. 2016; Wang et al. 2019). Expression of *Klrg1* was similarly localized to cells arising from the MCMV-ie2-gp33 infection, consistent with the known effector-memory phenotype of inflationary CD8+ T cells (Welten et al. 2020) (Figures 2B, S2D).

**Figure 2.**
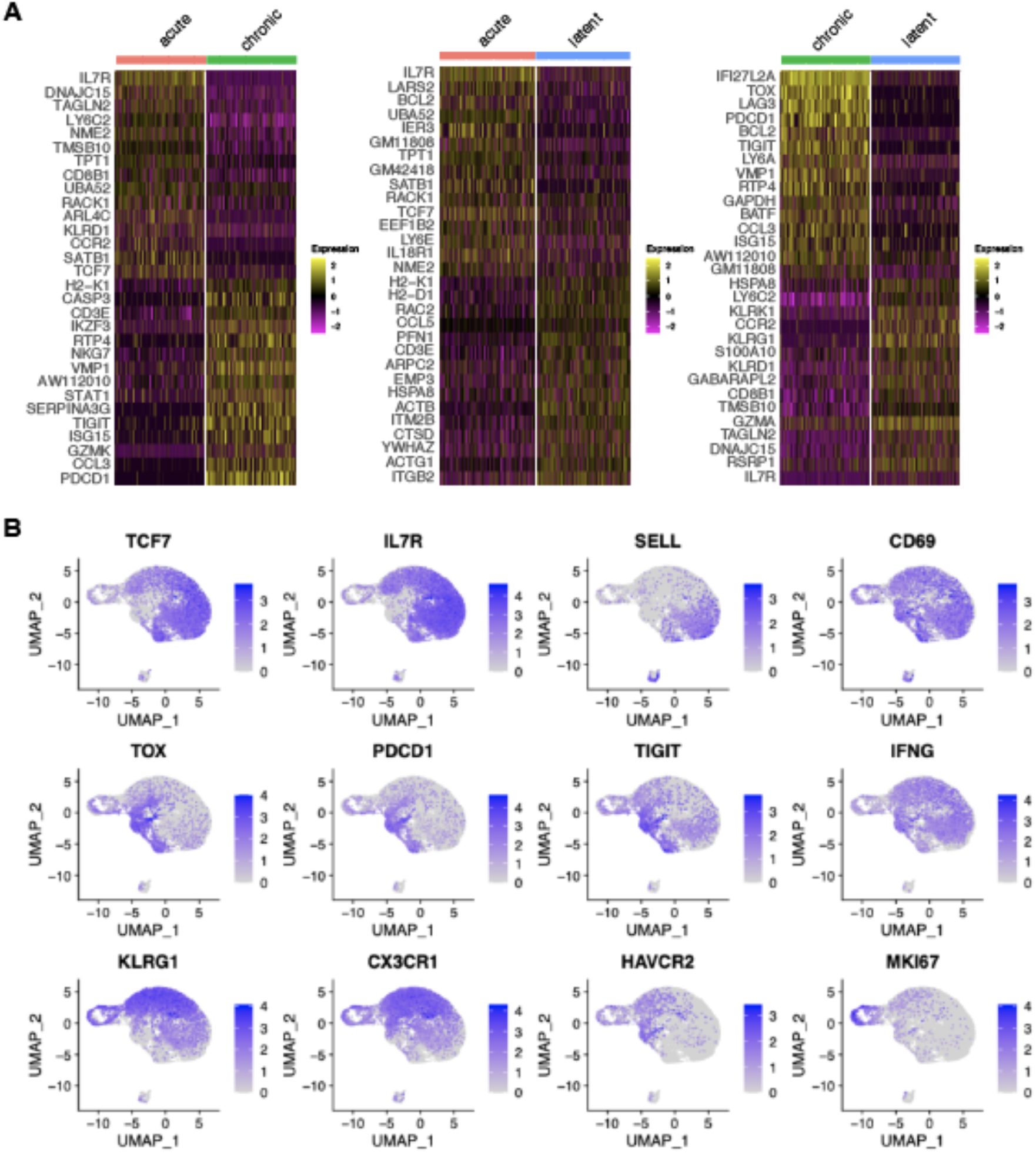
Virus-specific T cells have infection-specific phenotypes. A. Differentially expressed genes between acute and chronic LCMV infection (left), acute and MCMV-ie2-gp33 infections (middle), and chronic and MCMV-ie2-gp33 (right) infections. The upper 15 genes (from top to bottom) correspond to the highest positive average log fold change. Genes 16-30 represent those genes with the lowest average log fold change. All genes displayed have adjusted p values < 0.01. B. Normalized expression for select differentially expressed genes. Each uniform manifold approximation (UMAP) contains cells from all mice.

To obtain an additional unbiased confirmation of the various virus-specific CD8+ T populations, we used the 100 most upregulated genes from our previous differential gene analysis (Figure 2A) to match gene signatures with gene ontology (GO) terms and pathway analysis, confirming distinct CD8+ T cell phenotypes for the three infections (Figures S4, S5). Finally, we questioned whether these differentially expressed genes were in agreement with previously published gene sets, in contrast to our handpicked selection of phenotypic markers (Figures S6, Table S1). Together, this further highlighted distinct transcriptional phenotypes arising from acute, chronic and latent infections.

### Polyclonal but individualized CD8+ T cell clonal expansion following acute, chronic and latent infection

After observing the transcriptionally diverse gene expression profiles following the different infection types, we determined if TCR repertoires showed infection-specific features. By restricting our analysis to cells containing only one TRA and one TRB sequence, we obtained information ranging from hundreds to thousands of cells from each mouse (Figures 3A, S7A). Quantifying the number of unique clones [defined by unique complementarity determining region 3 beta (CDRb3) + CDR3 alpha (CDRa3) nucleotide sequence] revealed hundreds of clones for each mouse (Figures 3A, S7B), indicating both a polyclonal GP33-specific repertoire in all three infection conditions and the presence of clonal expansion. Next, we visualized the percentage of the repertoire comprised by each clone, which demonstrated that mice which had received acute LCMV infection (and therefore had already cleared the virus 28 dpi) had a higher fraction of clones supported by only a single cell barcode (Figure 3A). Quantifying the Shannon evenness, a commonly used entropy metric that provides a global view of clonal frequencies (Miho et al. 2018; Yermanos, Kräutler, et al. 2020; Greiff et al. 2017), further confirmed the notion that acute LCMV infection resulted in relatively less clonal expansion than the other two infection types, where antigen is still present (Figure S7C). A closer examination of the 30 most expanded clones from each repertoire (Figure 3A) revealed that in many cases individual clones were represented by hundreds of cells, particularly in chronically and latently infected mice, and, in the case of a single latently infected mouse, more than a thousand cells (Figure 3B).

**Figure 3.**
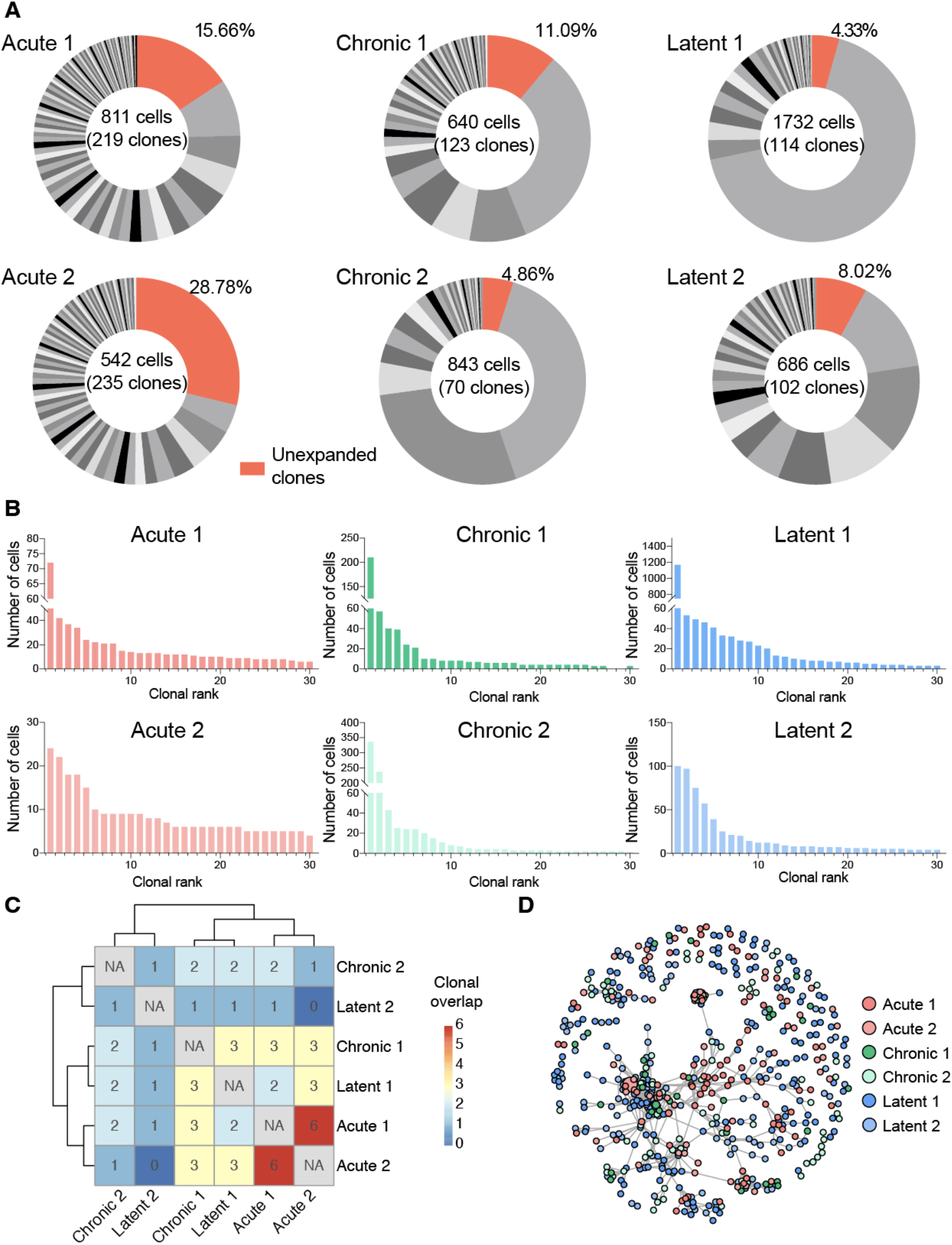
Virus-specific T cells are clonally expanded and personalized. A. Distribution of clonal expansion. Clone was defined by identical CDRb3-CDRa3 nucleotide sequence. Each section corresponds to a unique clone and the size corresponds to the fraction of cells relative to the total repertoire. Unexpanded clones were those clones supported by only one unique cell barcode. B. Clonal frequency for the top 30 most expanded clones in each repertoire. C. Number of identical clones found between mice. D. Similarity network of virus-specific CD8+ T cell clones. Nodes represent a unique CDRb3-CDRa3 from each mouse. Edges connect those clones separated by an edit distance of 5 amino acids or less.

We next determined the extent of clonal convergence in the GP33-specific repertoire by quantifying the number of identical TCRs found in each mouse, which revealed minimal overlap detected regardless of infection condition (Figure 3D). We observed no prominent relationship between clonal expansion and clonal overlap, as performing a similar analysis restricted to expanded clones (clones supported by two or more cells), including the 10 most expanded clones per mouse, did not reveal any substantial overlap (Figures S7D, S7E). We subsequently questioned if signs of clonal convergence could be detected by focusing the analysis on clones with similar, but non-identical TCR sequences. We therefore constructed sequence similarity networks based on the edit distance of the CDRb3 and CDRa3 sequences. Despite investigating a range of edit distance thresholds, we visually failed to observe any infection-specific clustering (Figures 3D, S8). Formally quantifying the number of edges shared either within the same or across different infection groups supported our visual observation that infection-specific overlap was not present (Figure S8).

### Stereotypic germline gene usage following acute, chronic and latent viral infection

After observing the low degree of sequence similarity across all mice, we determined if stereotypical patterns of germline gene usage could be observed in any of the infection conditions, as previous repertoire sequencing experiments investigating virus-specific T cells in the context of LCMV or MCMV infection have demonstrated preferential germline use (Welten et al. 2020; Yermanos, Sandu, et al. 2020). Leveraging the ability of our single-cell immune repertoire profiling, we could investigate the variable (V) gene usage for both TRB and TRA chains, in addition to quantifying how often certain pairings occured. Quantifying and visualizing the number of cells using a given germline pairing revealed certain V genes dominated the repertoire across multiple mice in different infection conditions, such as TRBV13-1, TRBV19, and TRBV29 (Figures 4A, S9A). Calculating pairwise correlation coefficients for germline gene usage between all mice demonstrated that TRBV gene usage loosely clustered repertoires by infection type (Figure 4B), although this effect was not observed when both TRBV and TRAV genes were included into the calculation (Figure S9B). We lastly questioned whether including repertoires lacking specificity to GP33 could provide contrast for how similar the 6 repertoires were following acute, chronic, and latent infection. Including publicly available data from either naive PBMCs (10x genomics) or CNS-resident T cells (Yermanos, Neumeier, et al. 2021) demonstrated clear separation between the repertoires of GP33-specific T cells and naive T cells (Figure 4C), further supporting the notion that the GP33-specific repertoire has stereotypic germline gene usage irrespective of the infection condition.

**Figure 4.**
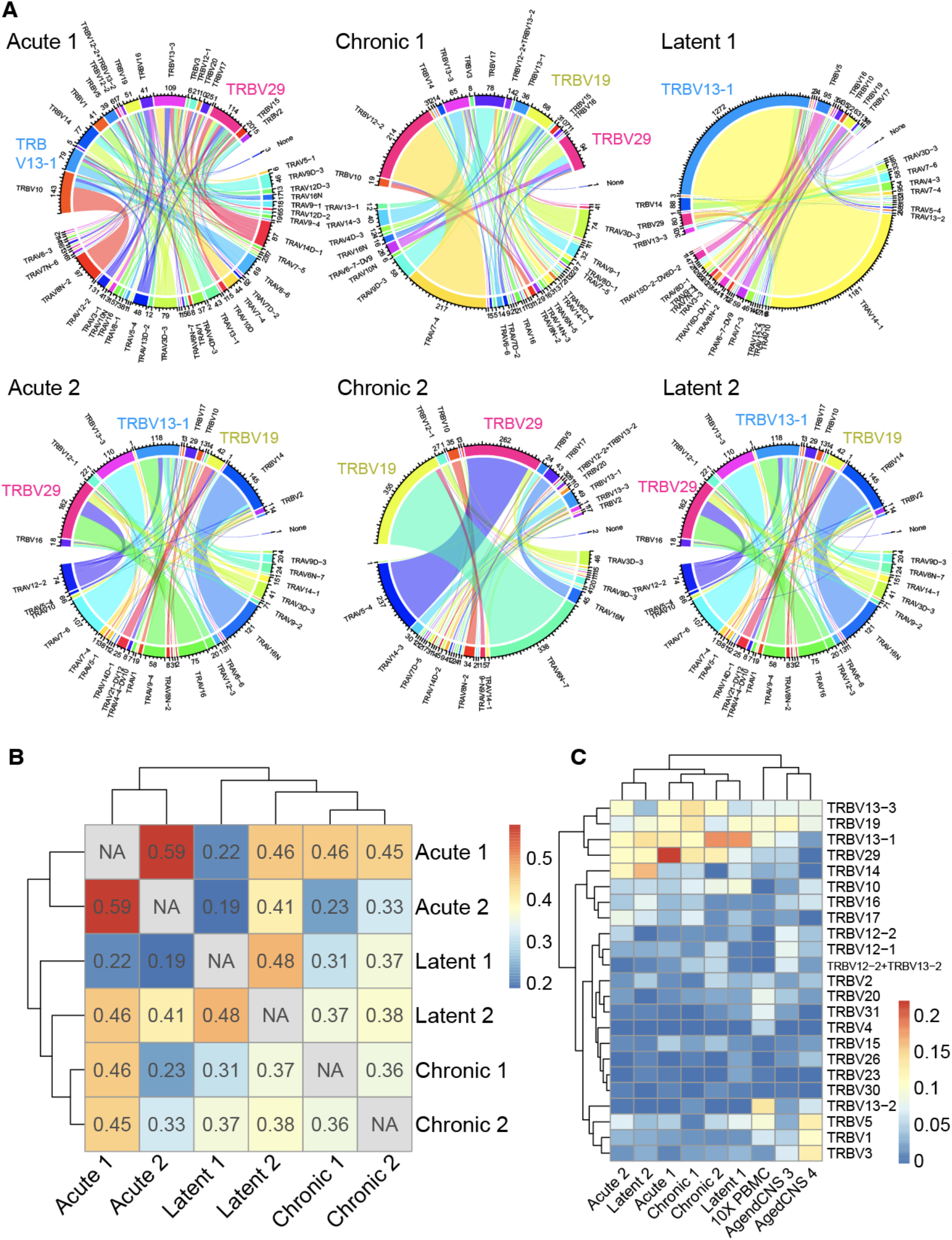
Stereotypic germline gene usage. A. Circos plots depicting the relationship between TRB and TRA V genes. Color corresponds to TRA gene usage. Connections illustrate the number of cells using each particular combination. B. Correlation heatmap quantifying the fraction of unique clones using a particular TRB V gene. Intensity corresponds to the Pearson correlation of the V gene usage vector between two samples. Clone was defined as an identical CDRb3-CDRa3 nucleotide sequence. C. TRB V gene usage compared to other single-cell immune repertoire sequencing datasets containing naive T cells.

### Transcriptional heterogeneity within expanded virus-specific clones

We next integrated TCR sequence with transcriptomes at single-cell resolution. It has been previously demonstrated that highly expanded clones upregulate effector molecules such as *Nkg7*, *Ccl5*, and granzymes (Yermanos, Agrafiotis, et al. 2021; Yermanos, Neumeier, et al. 2021). Therefore, we first focused our analysis on the 30 most expanded clones for each infection by quantifying the fraction of cells present in each transcriptional cluster (Figure 5A). Cells arising from different infection conditions occupied distinct transcriptional states, thereby suggesting transcriptional heterogeneity within the majority of expanded clones (Figure 5A). Extending this analysis to all clones, regardless of clonal expansion, would reveal differences between highly and lowly expanded clones. This analysis showed that more expanded clones (thicker lines in Figure 5B) were predominantly connected to clusters 5 and 2 for chronic and latent infections, respectively, whereas unexpanded clones (narrower lines) were often connected to clusters 7 and 0, respectively (Figures 5B, S10). Quantifying the cluster membership demonstrated a clear trend consistent in all six mice that clones with higher expansion were located in cluster 1 for acute, 5 for chronic, 2 for latent infection (Figure 5C), which characteristically expressed *Zeb2*, *Tox*, and *Klrg1*, respectively (Figures 1B, 1F, S2). Conversely, unexpanded clones were more often located in clusters 4 and 7 (Figure 5C), which were characterized by high expression of genes associated with memory phenotypes such as *Id3*, *Sell*, and *Tcf1* (Figures 1B, 1F, S2). Calculating the differentially expressed genes revealed that genes such as *Nkg7*, *Lgals1*, *Pdcd1*, and *Ccr2* were significantly upregulated in expanded cells in at least one infection condition and demonstrated consistent trends in expression for all infection groups (Figures 5D, S11A, S11B). Taken together, our findings support previously proposed models in which clonally expanded T cells remain transcriptionally heterogeneous yet preferentially adopt effector-like phenotypes.

**Figure 5.**
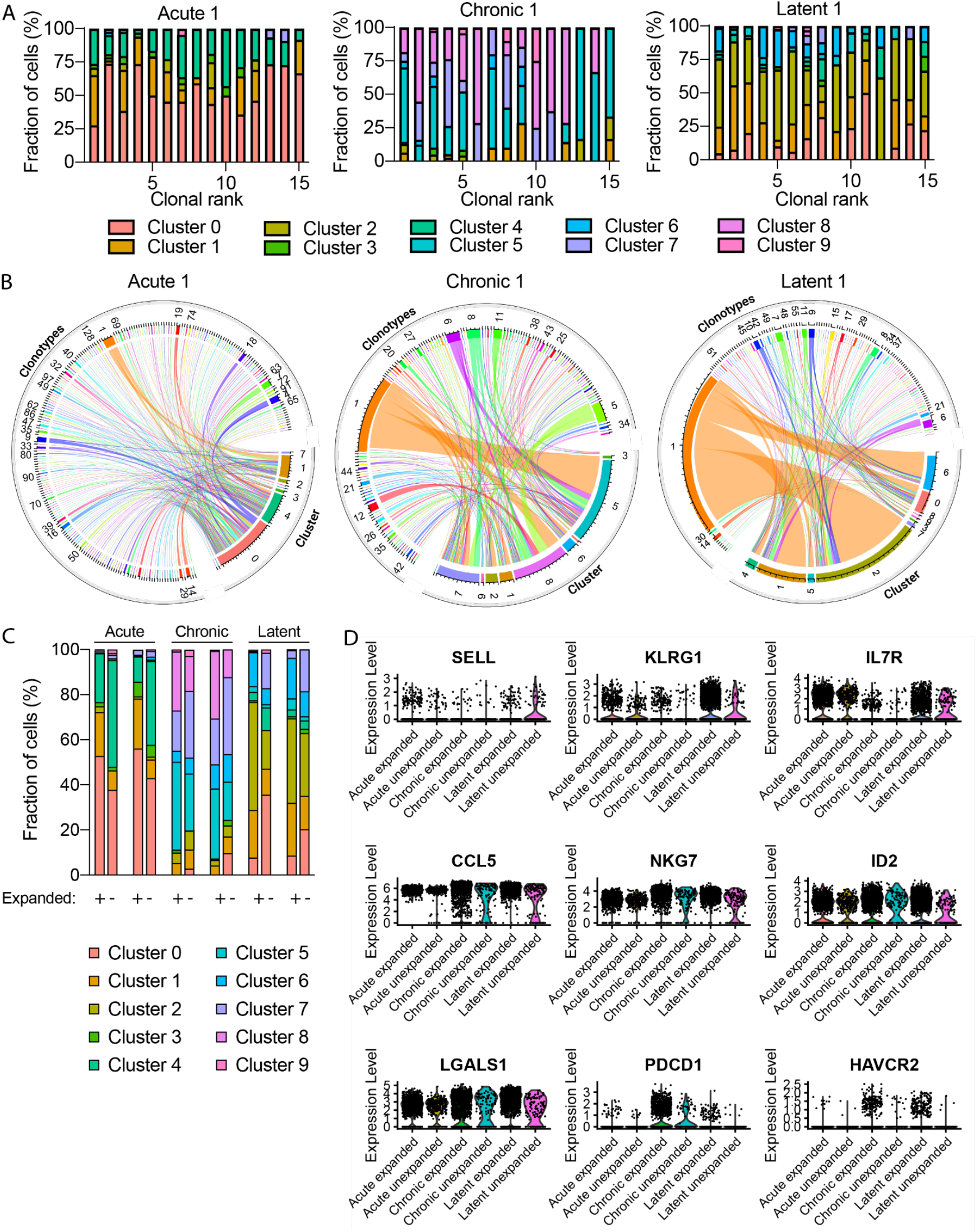
Transcriptional phenotypes relate to clonal expansion. A. Transcriptional cluster membership for the top 15 most expanded clones for each infection group. Clone was defined as identical CDRb3-CDRa3 nucleotide sequence. B. Circos plots relating cluster membership to clonal expansion. Numbers surrounding the circle indicate the number of cells within the specific clone. C. Transcriptional cluster membership for expanded (+) and unexpanded (-) clones for each mouse. Unexpanded clones were those clones supported by only one unique cell barcode. D. Violin plots for differentially expressed genes between expanded vs unexpanded cells for each infection condition.

### Pseudotime analysis supports clonal heterogeneity

Until now, our single-cell analysis has investigated transcriptional signatures after integrating all cells from the three infection types. We therefore next questioned whether first separating the cells by infection type and leveraging pseudotime analyses would provide additional insight into differentiation of repertoire and transcriptome features underlying clonal selection. We therefore realigned raw sequencing reads to the reference transcriptome and supplied the subsequent alignment files to the DropEst and Velocyto pipelines (La Manno et al. 2018; Petukhov et al. 2018). Performing unsupervised Louvain clustering and UMAP on the output of DropEst and incorporating vector fields from Velocyto highlighted transcriptional heterogeneity within each infection, with phenotypes corresponding to memory, effector, exhausted, and inflationary T cells (Figures 6A, S12, S13, S14). As Velocyto results in a vector field indicating cell-state differentiation by comparing ratios of unspliced to spliced RNA (La Manno et al. 2018), we questioned whether overlaying the most expanded clones would reveal interclonal differentiation trajectories. Visualizing both the Velocyto vector fields and the most expanded clones from each mouse revealed clonal heterogeneity in regards to location on the UMAP and transcriptional cluster, however, this suggested that clear intraclonal trajectories could not be resolved outside of the clusters corresponding to proliferation (Figures 6B, S15).

**Figure 6.**
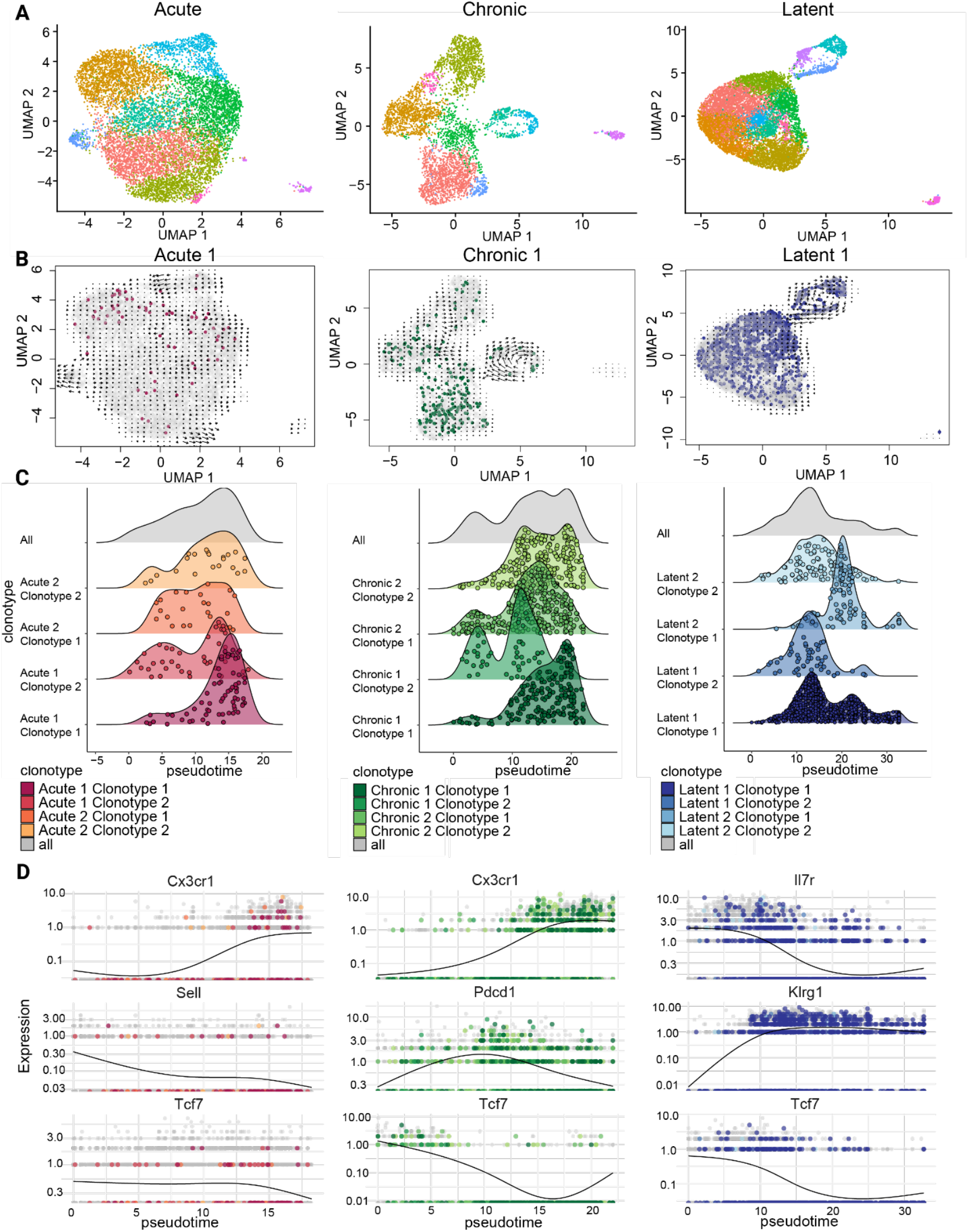
Pseudotime inference supports transcriptional heterogeneity for expanded virus-specific clones A. Uniform manifold approximation projection (UMAP) performed separately for virus-specific T cells from acute, chronic, and latent infections. Each point represents a cell and color indicates transcriptional cluster, unique to each UMAP. UMAP was created using DropEst and Velocyto. B. Pseudotime vector fields for each of the infection conditions. Points correspond to the most expanded clone found in each infection type. Clone was defined by identical CDRb3-CDRa3 nucleotide sequence. C. Monocle-inferred distribution of pseudotime for the most expanded clones in each mouse. Each point represents a cell. D. Monocle-inferred pseudotime for select genes for the most expanded clones in each mouse. Each point represents a cell.

Despite the inability of Velocyto to infer clear trajectories for all three infections, we questioned whether another method of calculating pseudotime, Monocle, could provide additional information regarding the differentiation of the most expanded clones. We therefore quantified the pseudotime state for the most expanded clones from each mouse and compared this to the remainder of cells from the respective repertoires, which indicated that the most expanded clones were distributed throughout multiple pseudotime states (Figures 6C, S16). After profiling the clonotype-specific pseudotime calculated using all genes, we restricted our analysis to specific genes of interest to compare the pseudotime heterogeneity with previously suggested differentiation states for acute, chronic, and latent infection (Zander et al. 2019; Hudson et al. 2019; Beltra et al. 2020). For the most expanded clones arising from acute LCMV infection, we observed a clear movement in pseudotime from a more memory-like phenotype (*Sell*) to an effector-like phenotype (*Cx3cr1*) (Figure 6D), whereas expanded clones from chronic LCMV infection resulted in a trajectory involving initial expression of *Tcf7*, then coexpression of *Tcf7* and *Pdcd1*, with cells of a later pseudotime expressing *Cx3cr1* (Figure 6D). In the case of MCMV infection, we observed that expanded clones occupied distinct states in pseudotime defined by expression of *Il7r*, *Klrg1*, and *Tcf7* (Figure 6D). Taken together, pseudotime analysis supported transcriptional heterogeneity of expanded clones yet also failed to clearly elucidate distinct differentiation pathways when based on mRNA splicing from Velocyto. Although this may be specific for our particular experimental setup, it is possible that either time-resolved scSeq data or entirely distinct experimental setups would result in higher resolution trajectories capable of quantifying differentiation at the single-clone resolution. We therefore developed a publicly available bioinformatic pipeline, Clonotyme, to integrate repertoire features with gene-expression based pseudotime inference (Figure S17). This workflow can take either the resulting alignment necessary for RNA velocity or an already computed Seurat object and will integrate clonal information to either method of pseudotime analysis. Upon specifying clonal features (e.g., expansion rank), the user can visualize individual clones in pseudotime dimensions, both relative to the global transcriptional landscape (e.g., directly on the UMAP) or for individual genes. Together, Clonotyme supported a model in which clonally-related virus-specific CD8+ T cells were located across diverse regions of pseudotime and can additionally expedite future time-resolved scIRS studies.

## Discussion

Here, we used single-cell TCR repertoire and transcriptome sequencing to investigate how T cell clonal selection signatures vary across acute, chronic, and latent viral infection in mice. While the recovered CD8+ T cells shared specificity to a common viral peptide, our results demonstrated infection-specific transcriptional heterogeneity that was maintained across biological replicates (Figure 1B). While previous reports have demonstrated that acute, chronic, and latent infections result in T cells with a range of phenotypes and effector functions (Wherry 2011; Alfei et al. 2019; Zehn et al. 2012; Utzschneider et al. 2016; Barnstorf et al. 2019; Hudson et al. 2019; Welten et al. 2020), a comparison characterizing whole transcriptomes at the single-cell level has not yet been performed. Our findings showcase the extensive T cell phenotypic diversity and similarly highlight the lack of transcriptional overlap between CD8+ T cell phenotypes from the three models of infection. Consistent with previous results, we could recreate the effector, memory, exhausted, and inflationary expression signatures characteristic of acute, chronic and latent infection using both targeted and unbiased computational analyses.

Previous experiments characterizing the endogenous GP33-specific TCR repertoire in the context of LCMV infection have demonstrated varying degrees of polyclonality (Yermanos, Sandu, et al. 2020; Chang et al. 2020). Leveraging TRB repertoire sequencing, both studies recovered multiple distinct clones, ranging from 40 to hundreds of unique GP33-specific CD8+ T cell clones following chronic and acute LCMV infection. The number of unique clones reported by both studies were comparable to the number of GP33-specific CD8+ T cells found in naive CD8+ T cells of uninfected C57BL/6 mice (Malhotra et al. 2020; Obar, Khanna, and Lefrançois 2008; Kotturi et al. 2008). Importantly, both TRB studies demonstrated high clonal overlap between the TCF1+ and TCF1-CD8+ T cell repertoires (Yermanos, Sandu, et al. 2020; Chang et al. 2020), which together supports a previously proposed model in which TCF1+ CD8+ T cells feed into the TCF1-CD8+ T cell subset (Utzschneider et al. 2016; Siddiqui et al. 2019). Similarly, a high degree of clonal overlap between the TCF1+ and TCF1-repertoires was observed in the context of inflationary T cells following MCMV infection (Welten et al. 2020). However, as these studies relied upon bulk TRB chain sequencing, relating clonality to gene expression profiles was not possible.

Our single-cell sequencing approach allowed us to relate individual transcriptomes to the TCR repertoire for thousands of cells, thereby providing insight into the relationship between gene expression and clonality. While we could again confirm a polyclonal response through detecting hundreds of unique GP33-specific clones following acute, chronic, and latent infections, we could, for the first time, demonstrate a polyclonal and expanded GP33-specific TCR repertoire at the single-cell resolution. We additionally discovered transcriptional diversity within individual clones that was present in each infection condition. Here, we again observed that clonally expanded T cells are found in both *Tcf1*+ and *Tcf1*- clusters, thereby supporting the previously reported model which implies a clonal relationship between the TCF1+ and TCF1- T cells during chronic viral infection (Utzschneider et al. 2016). While this hypothesis has been similarly described in the context of cancer (Siddiqui et al. 2019) and MCMV infection (Welten et al. 2020), an extensive characterization of this hypothesis at the polyclonal GP33-specific repertoire level was lacking.

In contrast to our previous findings (Yermanos, Sandu, et al. 2020; Welten et al. 2020), the virus-specific CD8+ TCR repertoires were extremely personalized, with minor clonal overlap between mice. This was true for both expanded and unexpanded clones, suggesting a stochasticity underlying the selection and expansion of virus-specific clones. The findings presented here may contrast to higher clonal overlap previously reported due to inherent differences in the repertoire sequencing technologies. Specifically, the 10x genomics platform used in this study provides unique molecular identifiers to reduce PCR and sequencing errors and additionally does not rely on multiplex primers, which should improve the accuracy and reduce amplification biases. Although we did not observe a high degree of clonal overlap, we observed that certain germline genes were used more often in TCRs with a common specificity to a single, shared viral epitope. As the naive repertoires of these mice are generated from identical TCR loci, our findings imply that the inflammatory context of distinct infection does not dictate the germline gene selection and accompanying preferential expansion as much as the exact specificity does.

## Methods

### Animal experiments

All animal experiments were performed in accordance with institutional guidelines and Swiss federal regulations. Experiments were approved by the veterinary office of the canton of Zurich under animal experimentation licenses 115/2017 and ZH058/20. 6-8 week old female C57BL/6 mice from Janvier were housed under specific-pathogen-free conditions in individually ventilated cages with bedding and nesting material for enrichment. Acute LCMV infections were infected intravenously (i.v.) with 200 focus forming units (ffu) of LCMV clone 13 in the tail vein. Chronic LCMV infections were performed i.v. with 2 x 10^6^ ffu LCMV clone 13. Latent infections were established by injecting 2×10^5^ pfu dose of MCMV-*ie2*-gp33 i.v., which was obtained from Dr. L. Cicin-Sain and contains a functional m157 gene as previously described (Welten et al. 2015). MCMV viral stocks were propagated on M2-10B4 cells and purified by ultracentrifugation using a 15% sucrose gradient. LCMV clone 13 waas produced as previously described (Sandu et al. 2020). Upon sacrifice with CO2 at 28 dpi, spleens were harvested and single-cell suspensions were prepared by mashing the tissue through a 70 uM cell strainer and rinsing with complete RPMI (RPMI-1640 supplemented with 10% fetal bovine serum, 2 mM L-glutamine, 1% penicillin-streptomycin, 1 mM sodium pyruvate, 50 nM beta-mercapthoethanol, 0.1 mM non-essential (glycine, L-alanine, L-asparagine, L-aspartic acid, L-glutamic acid, L-proline, L-serine) amino acids, 20 mM HEPES). The single cell suspension was then incubated with CD8-PE (clone 53-6.7, Biolegend), MHC class 1 tetramer for gp_33-41_ conjugated to APC diluted in FACS buffer (PBS, 2 mmEDTA, 2% FCS) at room temperature for 30 minutes, as previously described (Altman et al. 1996), and LiveDead nearIR. Tetramer positive cells were isolated via flow cytometric sorting (FACSAria with FACSDiva software) and subsequently supplied as input for single-cell immune repertoire sequencing.

### Single-cell immune repertoire sequencing

Single-cell immune repertoire sequencing was performed as according to the 10x Genomics Chromium Single Cell V(D)J Reagents Kit (CG000166 Rev A) as previously described (Neumeier et al. 2021). In brief, single cells for all six samples were simultaneously encapsulated with gel emulsion microdroplets (10x Genomics, 1000006) in droplets using 6 lanes of one Chromium Single Cell A Chip (10x Genomics, 1000009) with a target loading of 13,000 cells per reaction. cDNA amplification was performed using 14 cycles and subsequently split for downstream GEX and VDJ library preparation. GEX libraries were amplified using the Chromium Single Cell 5’ Library Kit (10x Genomics, 1000006). TCR libraries were amplified using the Chromium Single Cell V(D)J Enrichment Kit, Mouse T Cell (10x Genomics, 1000071). Final libraries were pooled and sequenced on the Illumina NovaSeq S1 using a concentration of 1.8 pM with 5% PhiX.

Paired-end sequencing files for GEX and VDJ libraries were aligned to the murine reference genome (mm10) and V(D)J germlines (GRCm38) using 10x Genomics cellranger (v4.0.0) count and vdj arguments, respectively. The filtered feature matrix directory was supplied as input to the automate_GEX function in the R package Platypus (v2.0.5) (Yermanos, Agrafiotis, et al. 2021), which uses the transcriptome analysis workflow of the R package Seurat (Satija et al. 2015). Only those cells containing less than 20% of mitochondrial reads were retained in the analysis. Genes involved in the adaptive immune receptor (e.g., TRB, TRBV1-1), were removed from the count matrix to prevent clonal relationships from influencing transcriptional phenotypes. Gene expression was normalized using the “scale.data” argument in automate_GEX, which first performs log-normalization with a scaling factor of 10000 and then scales mean expression and variance to 0 and 1, respectively. 2000 variable features were selected using the “vst” selection method and used as input to principal component analysis (PCA) using the first 10 dimensions. Graph-based clustering using the Louvain modularity optimization and hierarchical clustering was performed using the functions FindNeighbors and FindClusters in Seurat using the first ten dimensions and a cluster resolution of 0.5. UMAP was similarly inferred using the first ten dimensions. The FindMarkers function from Seurat was used when calculating differentially expressed genes (both across groups or across clusters) with both the minimum log fold change and the minimum number of cells expressing each gene set to 0.25. Mitochondrial and ribosomal genes were removed when either visualizing DE genes or supplying the top DE genes as input to gene ontology and gene set enrichment analyses. Gene ontology and gene set enrichment analysis was performed using the GEX_GOterm and GEX_GSEA functions in Platypus by supplying either the top N or bottom N genes as input. In the case of GEX_GSEA, the C7 immunological signatures gene set from the Broad institute was supplied as input to the function (Subramanian et al. 2005). The GEX_GSEA function uses the R package fgsea (Korotkevich, Sukhov, and Sergushichev 2019) to conduct gene set enrichment analysis and GEX_GOterm is based on the R package edgeR (Robinson, McCarthy, and Smyth 2010).

For TCR repertoire analysis, the output directory of 10x Genomics cellranger vdj function was supplied as input to the VDJ_analyze function in Platypus maintaining the default clonotyping strategy (CDRa3+CDRb3 nucleotide sequence) as performed by cellranger. Those clones not containing exactly one TRA and one TRB chain were removed from the analysis. Clonal frequency was determined by counting the number of distinct cell barcodes for each unique CDR3. Overlap matrices were calculated by first appending the CDRa3 and CDRb3 nucleotide sequences and then quantifying the exact matches across repertoires. Similarity networks were calculated based on the VDJ_network function in Platypus, which first calculates the edit distance separately for TRB and TRA CDR3s, and then draws edges between those clones with a distance below the specified threshold. Circos plots were created using the VDJ_circos function in Platypus with a label.threshold of 5. Those cells in clones supported by only one cell were considered unexpanded clones, whereas those clones supported by two or more cells were considered expanded. Pseudotime analysis was performed by aligning the output bam files from cellranger using DropEst (Petukhov et al., 2018). Cells with nCount_spliced less than 1000 and more than 20% mitochondrial genes, as well as genes involved in the adaptive immune receptor were filtered out and the object loaded as a Seurat object analogously to the automate_GEX() funcion using “scale.data” as a normalisation method for gene expression, log-normalisation with a scaling factor of 10000, 2000 selected variable features using the “vst” selection method and the first 10 dimensions used as input to principal component analysis (PCA) and UMAP. Clustering was conducted as described previously. Subsequently velocyto.R (Kharchenko 2018) was run using the RunVelocity function from the SeuratWrappersR package (Satija et al. 2020). Following parameters were used: deltaT = 1, kCells = 25 and fit.quantile = 0.02. The resulting Seurat object was transformed to a Cell Data Set using the as.cell_data_set function from the SeuratWrappers package and was subsequently supplied as input to Monocle3 to calculate infection specific trajectories (Trapnell et al. 2014, Qiu et al. 2017, Cao et al. 2019, McInnes 2018), with the clusters containing central memory genes (2 for acute, 2 for chronic, and 2 for latent) set as the root of the trajectories. Clusters containing B cells (cluster seven for acute, seven for chronic, and ten for latent) were excluded from this analysis. The density distribution of the two most expanded Clonotypes for each sample across monocle-based pseudotime was visualized using the R package ggridges. The expression level of selected genes across monocle-based pseudotime was inferred using the plot_genes_in_pseudotime function of Monocle 3.

#### Data visualization

Heatmaps displaying differential gene expression were produced using the DoHeatmap function in the R package Seurat (v4.0.1) (Butler et al. 2018). Gene enrichment analysis was performed using the GEX_GOterm function in Platypus, which is based on the analysis pipeline in edgeR (Robinson, McCarthy, and Smyth 2010). Enrichment plots were produced using the R package ggplot (Wickham and Wickham 2007). Gene set enrichment analysis was performed using the GEX_GSEA function in Platypus (v3.1) under default parameters, which utilizes fgsea (v1.12), tibble (v2.1.3), and the C7 gene set from the molecular signatures database MSigDB (Korotkevich, Sukhov, and Sergushichev 2019; Liberzon et al. 2015). Similarity networks were produced using the R package igraph (Csardi, Nepusz, and Others 2006). Circos plots were produced using the chordDiagram function of the R package Circlize (Gu, 2014). All other figures were produced using Prism v9 (Graphpad).

## Acknowledgements

We acknowledge and thank Dr. Christian Beisel, Elodie Burcklen, and Ina Nissen at the ETH Zurich D-BSSE Genomics Facility Basel for excellent support and assistance. We also thank Nathalie Oetiker and Franziska Wagen for excellent experimental support.

---

**Figure S1.**
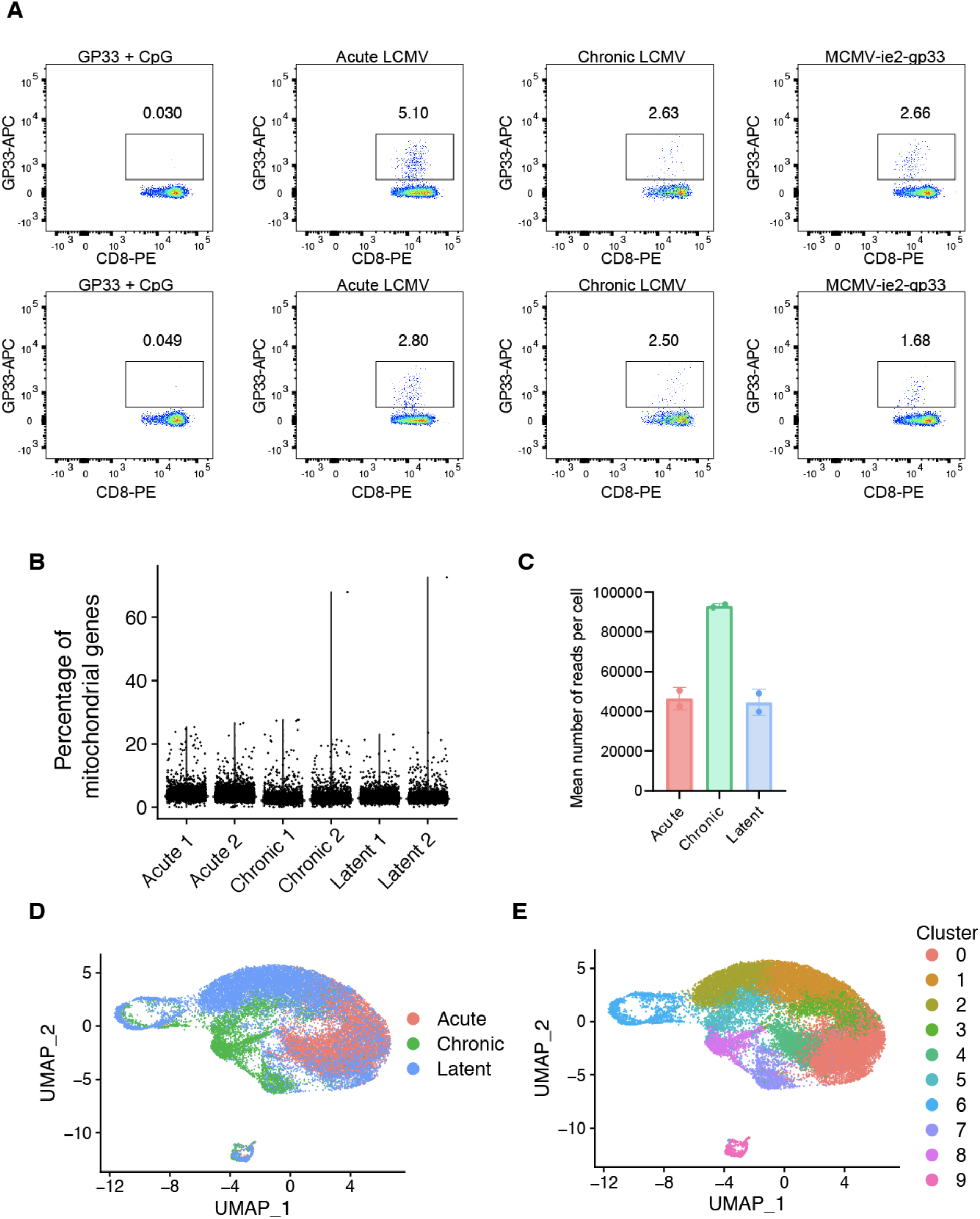
Single-cell sequencing of virus-specific CD8+ T cells. A. Flow cytometry plots of tetramer-sorted GP33-specific cells for all 6 mice. Pre-gated on lymphocytes, singlets, alive+ CD8+ cells. Due to the low number of cells, single-cell sequencing of the GP33 + CpG group was not performed. B. Percentage of mitochondrial genes per cell. C. Uniform manifold approximation projection (UMAP) colored by infection group. Each point represents a cell and color corresponds to transcriptional clusters. All cells from all samples were integrated in this single UMAP. D. Uniform manifold approximation projection (UMAP) colored by transcriptional cluster.

**Figure S2.**
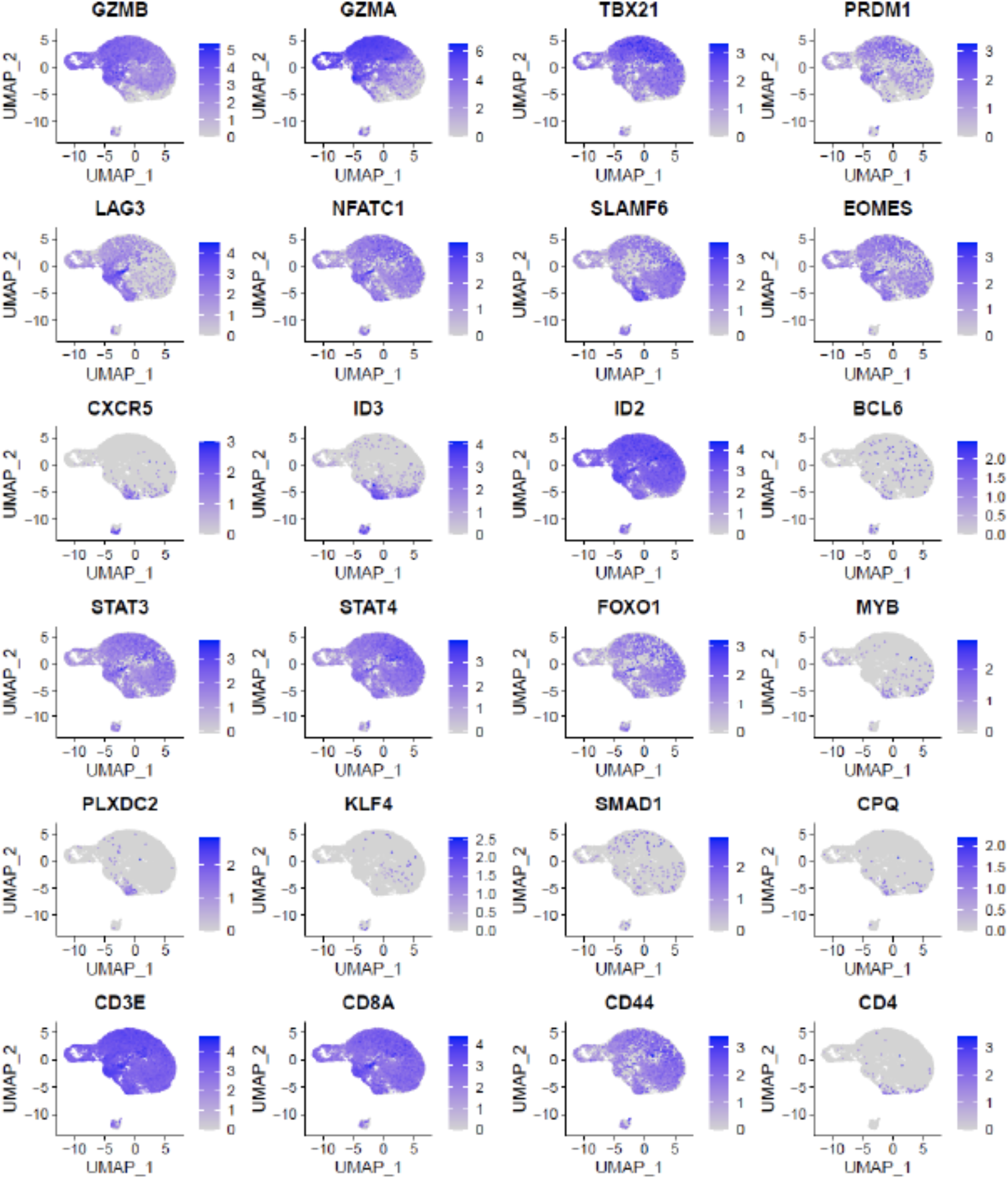
Normalized gene expression for select genes of interest. All cells from all samples were integrated into a single uniform manifold approximation projection (UMAP).

**Figure S3.**
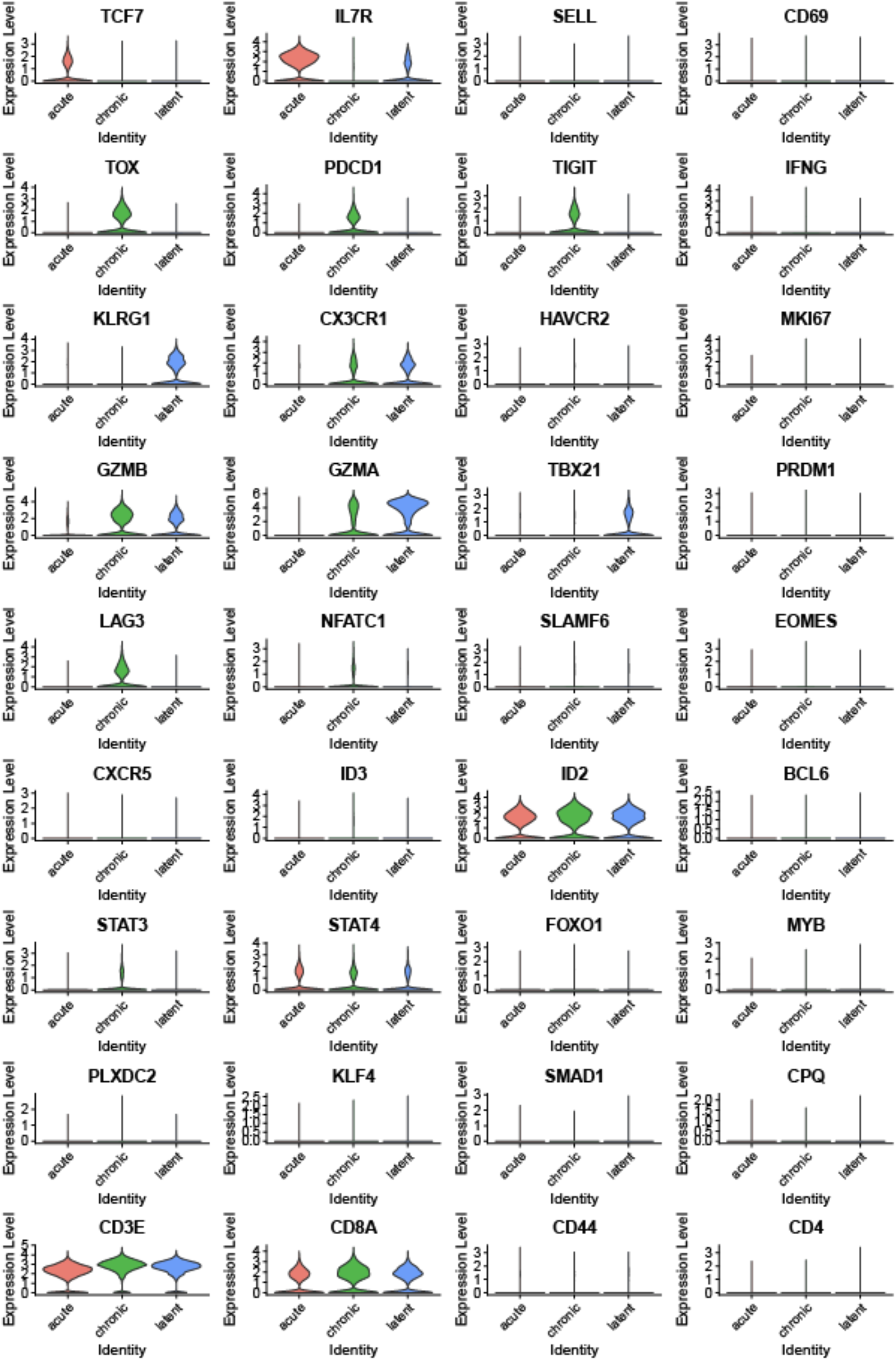
Normalized gene expression for select genes of interest split by infection type.

**Figure S4.**
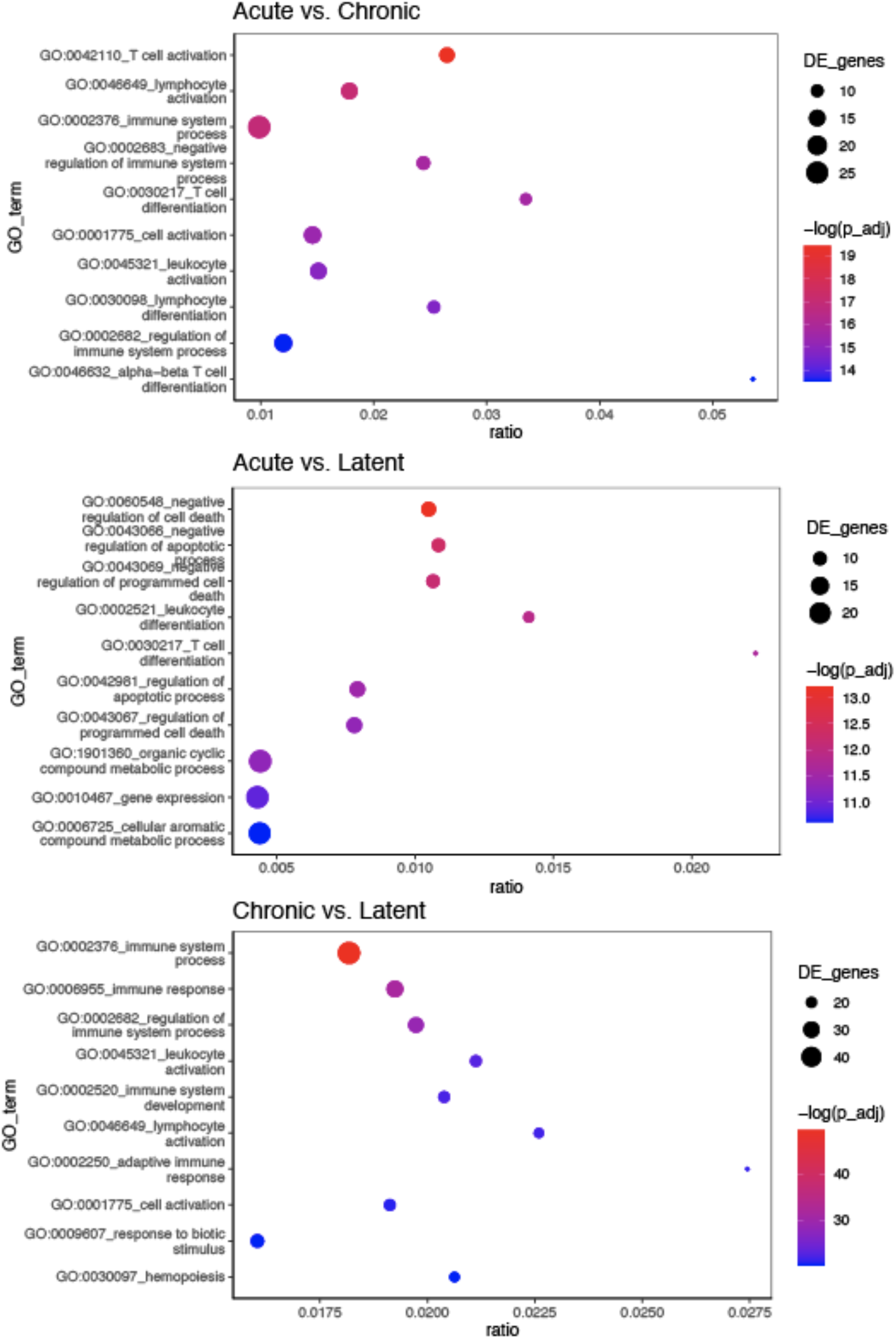
Gene ontology (GO) term enrichment of the 10 most upregulated genes from either acute versus chronic LCMV infection (top), acute LCMV versus MCMV-*ie2*-gp33 infection (middle), or chronic versus MCMV-*ie2*-gp33 (bottom) infection. The color of each dot corresponds to adjusted p value. The size of the dot corresponds to the number of genes. Ratio corresponds to the number of differentially genes relative to the number of total genes corresponding to each GO term.

**Figure S5.**
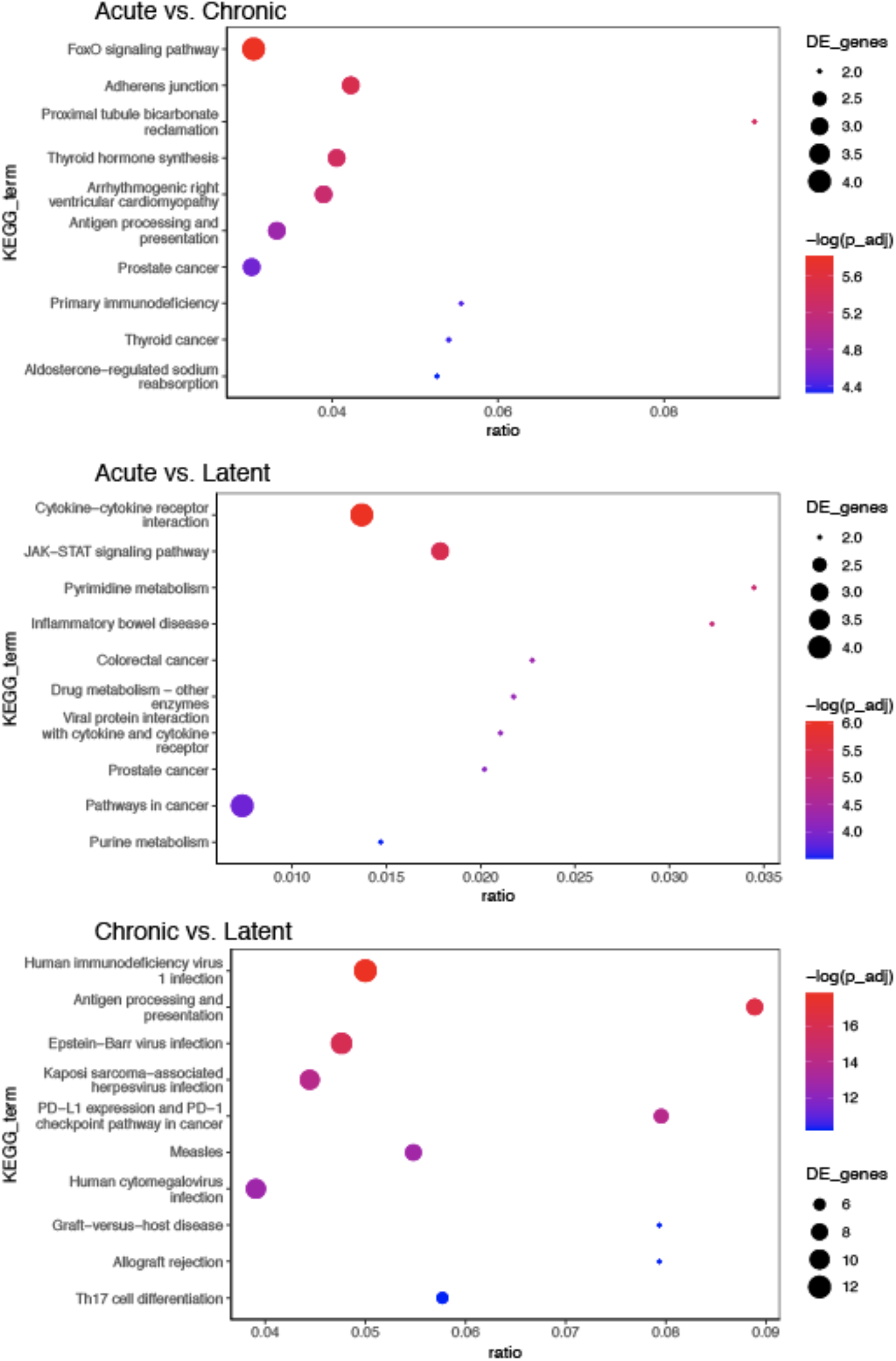
Pathway enrichment of the 10 most upregulated genes from either acute versus chronic LCMV infection (top), acute LCMV versus MCMV-*ie2*-gp33 infection (middle), or chronic versus MCMV-*ie2*-gp33 (bottom) infection. The color of each dot corresponds to the adjusted p value. The size of the dot corresponds to the number of genes. Ratio corresponds to the number of differentially genes relative to the number of total genes corresponding to each pathway.

**Figure S6.**
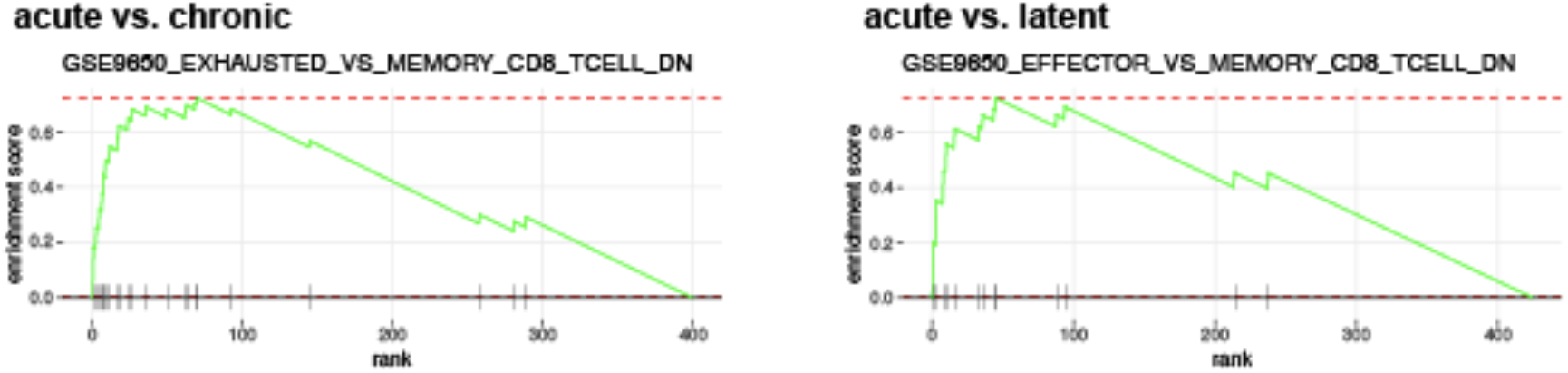
Gene set enrichment (GSEA) plots based on the C7 immunological signatures from the Broad institute. The upregulated genes from either acute versus chronic LCMV infection or acute versus MCMV-*ie2*-gp33 infection were supplied as input.

**Figure S7.**
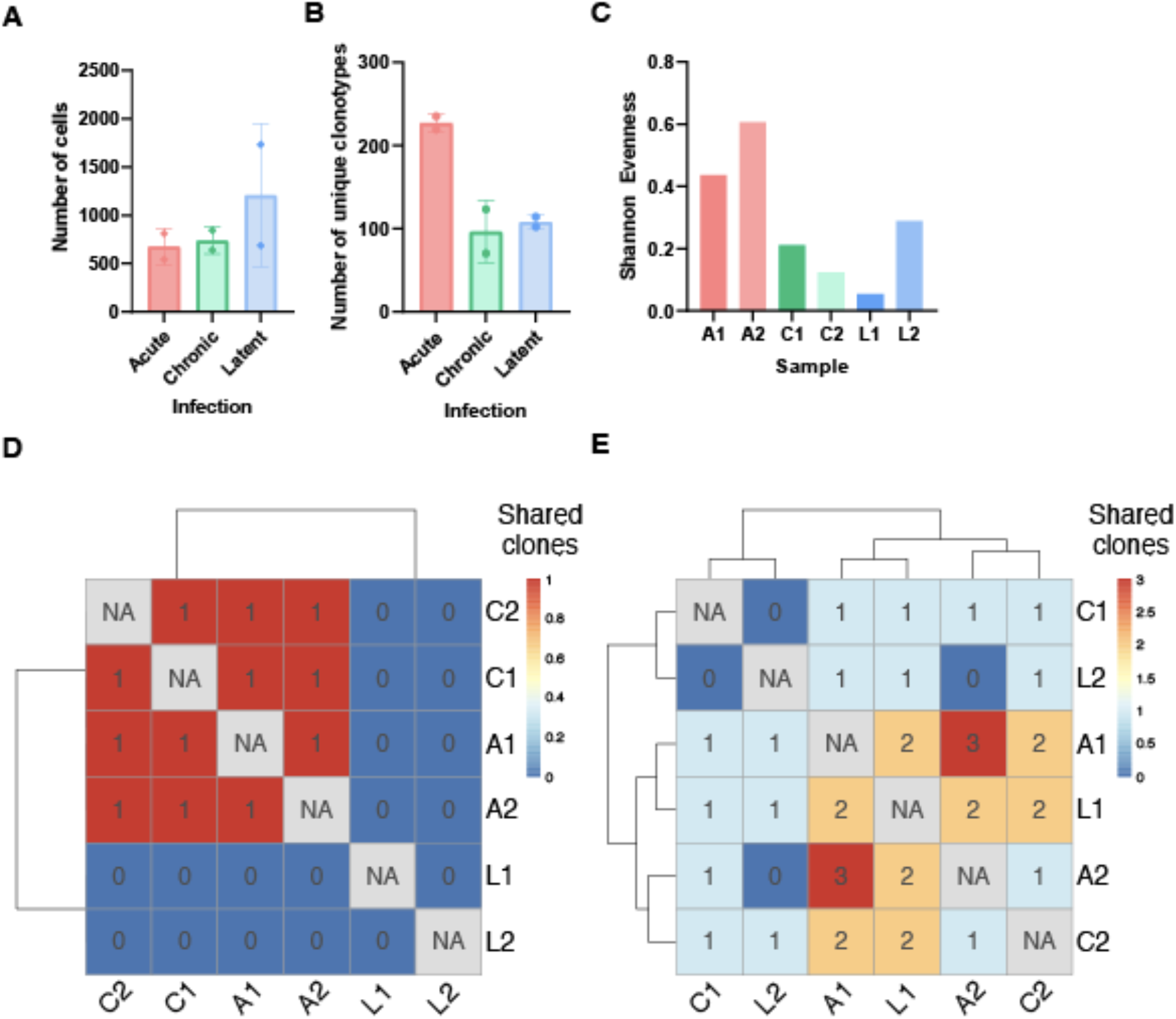
T cell receptor repertoire features of virus-specific CD8+ T cells following acute, chronic, and latent viral infection. **A.** Number of recovered GP33-specific cells containing exactly one T cell receptor beta (TRB) and T cell receptor alpha (TRA) B. Number of unique clones per repertoire. Clone was defined by identical CDRb3-CDRa3 nucleotide sequence. C. Shannon evenness quantifying the distribution of clonal frequency for mice infected with either acute (A1, A2), chronic (C1, C2), or latent (L1, L2) infection. D. Heatmap showing pairwise clonal overlap for the top 10 most expanded GP33-specific T cell clones from each mouse. E. Heatmap showing pairwise clonal overlap for those clones supported by two or more distinct cell barcodes.

**Figure S8.**
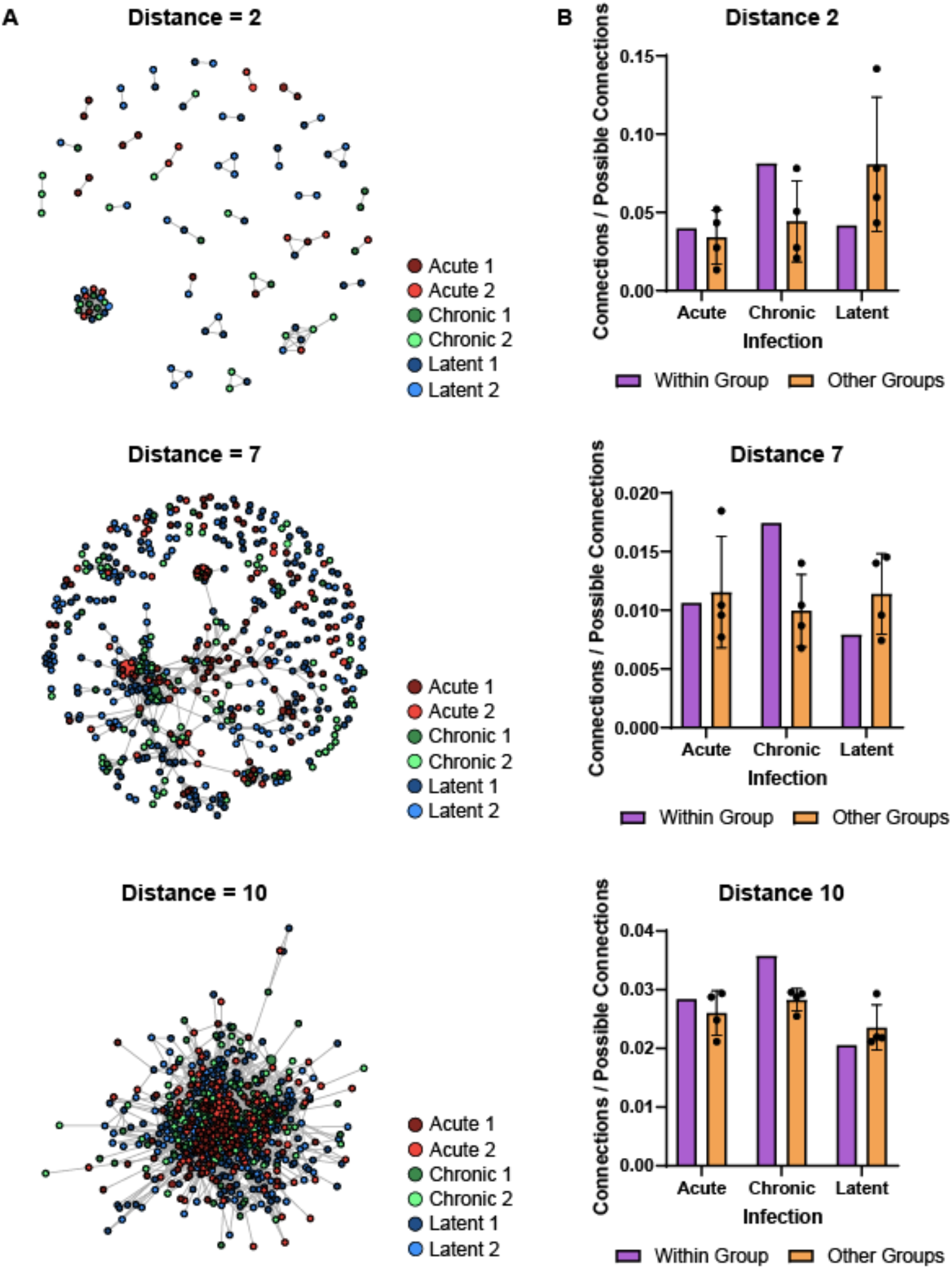
Similarity networks with diverse amino acid edit distance thresholds. A. Similarity network of virus-specific CD8+ T cell clones. Nodes represent a unique CDRb3-CDRa3 from each mouse. Edges connect those clones separated by an edit distance of N amino acids or less. B. The number of edges between nodes of mice either in the same infection group or with different infection groups normalized by the number of possible connections.

**Figure S9.**
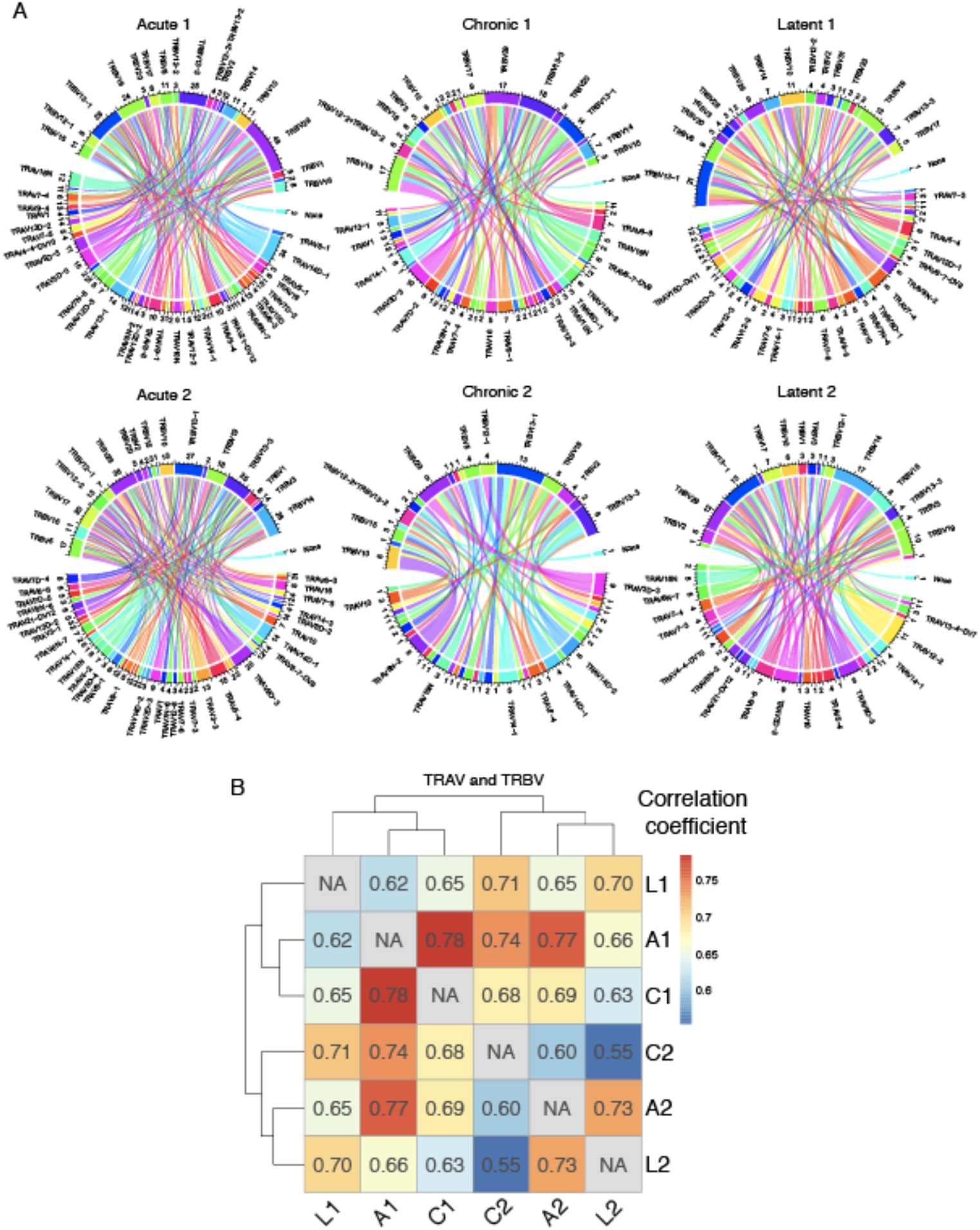
Germline gene usage of GP33-specific CD8+ T cell receptors. A. Circos plots depicting the relationship between TRB and TRA V gene usage. Color corresponds to TRA gene usage. Connections illustrate the number of clones using each particular combination. Clone was defined as an identical CDRb3-CDRa3 nucleotide sequence. B. Correlation heatmap quantifying the fraction of unique clones using a particular V gene. Intensity corresponds to the Pearson correlation of the V gene usage vector between two samples.

**Figure S10.**
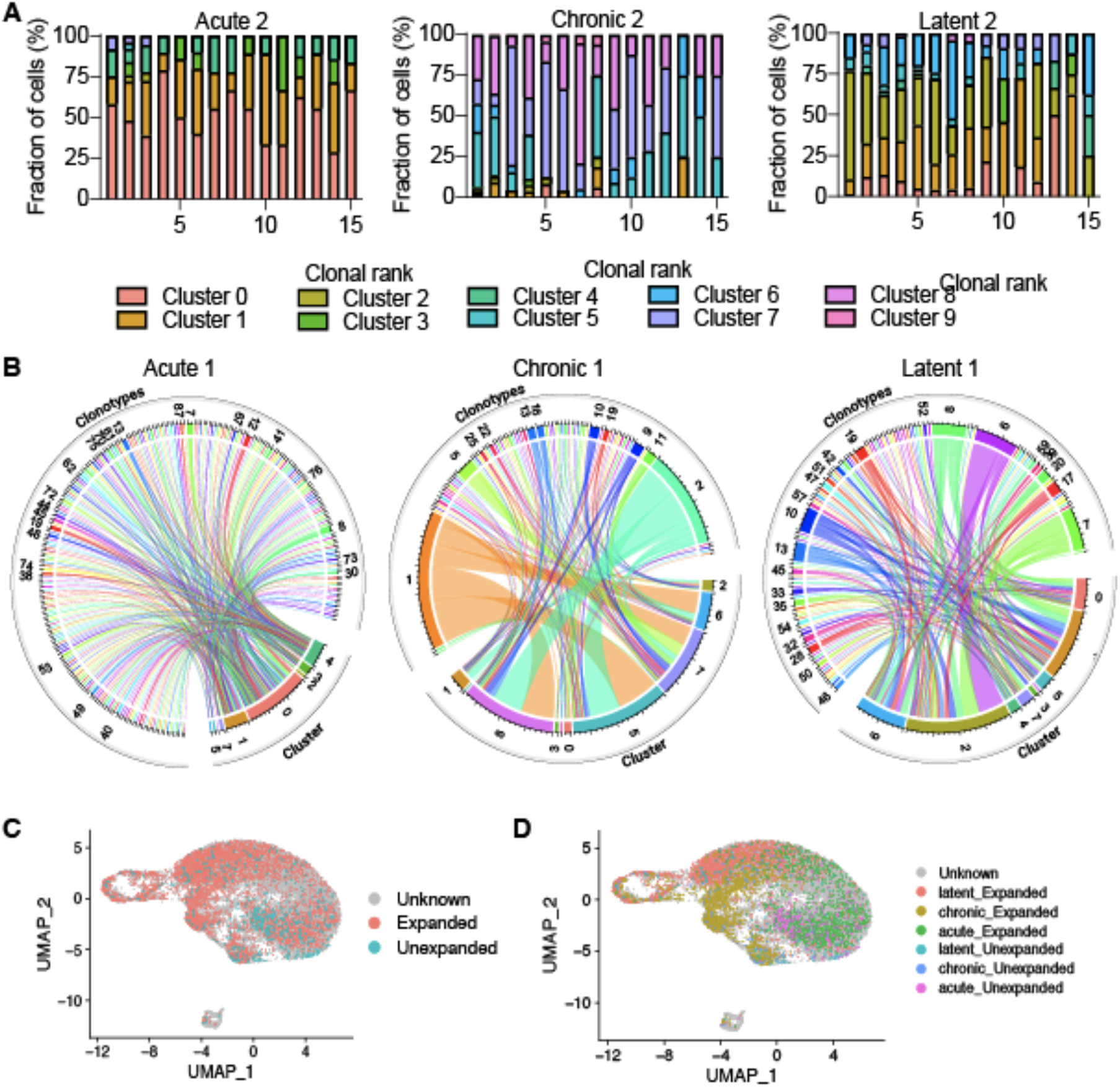
Supporting information for cluster membership. A. Transcriptional cluster membership for the top 15 most expanded clones for each infection group. Clones were defined as identical CDRb3-CDRa3 nucleotide sequence. B. Circos plots relating cluster membership to clonal expansion. Color corresponds to each unique clone. Connections illustrate the number of clones using each particular combination. Clone was defined as an identical CDRb3-CDRa3 nucleotide sequence. C. Uniform manifold approximation projection (UMAP) showing expanded (>1 cell) and unexpanded (1 cell) clones. Each point is a cell and cells from all infection conditions and mice were pooled. D. UMAP visualizing expanded and unexpanded clones but colored by infection type.

**Figure S11.**
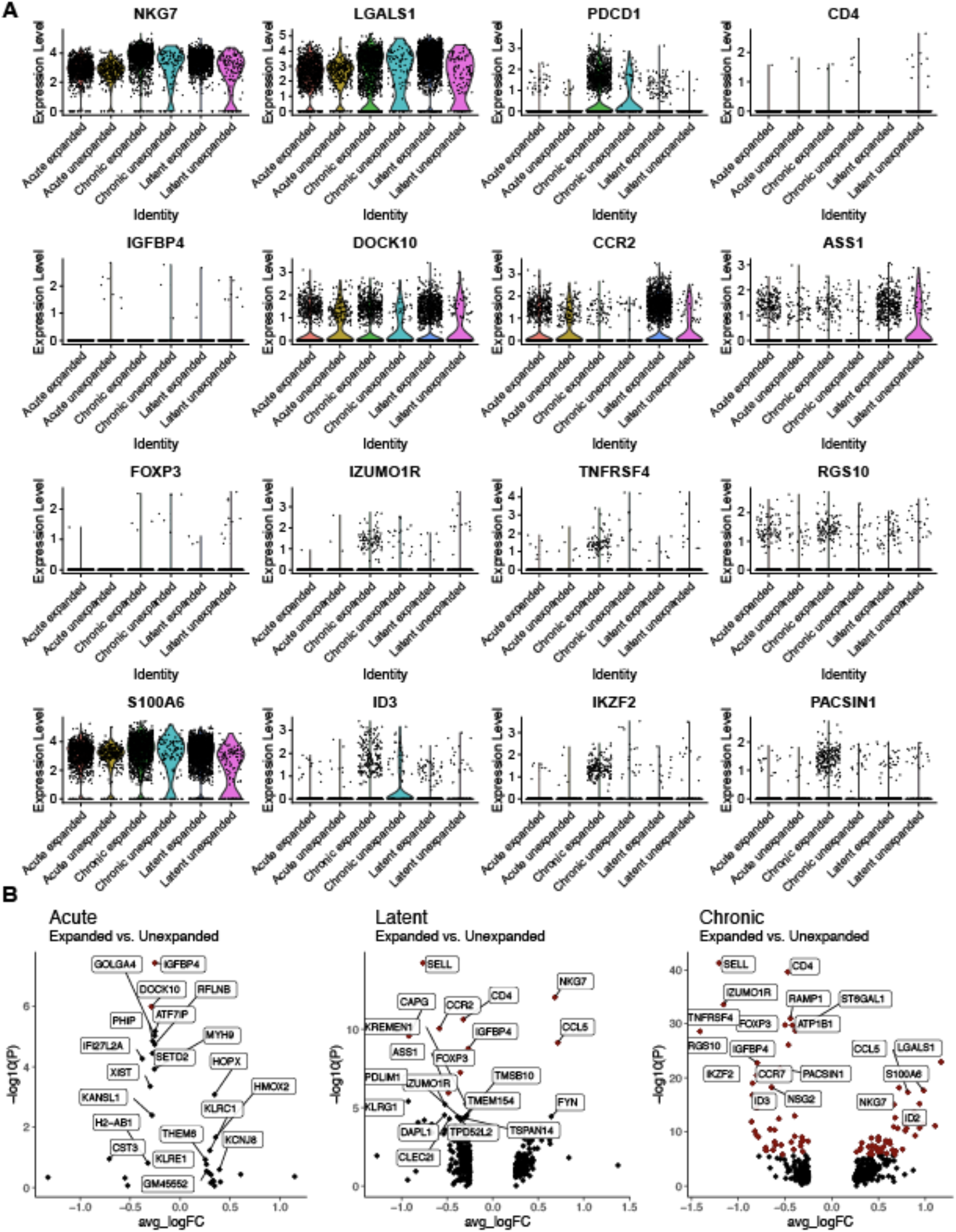
Expanded clones have distinct transcriptional profiles. A. Normalized gene expression for select genes of interest split by infection type and clonal expansion. Clone was defined as an identical CDRb3-CDRa3 nucleotide sequence. Expanded refers to those clones supported by more than one unique cell barcode. B. Differential gene expression between expanded and unexpanded cells in the three infection conditions. Points in red indicate significantly differentially expressed genes.

**Figure S12.**
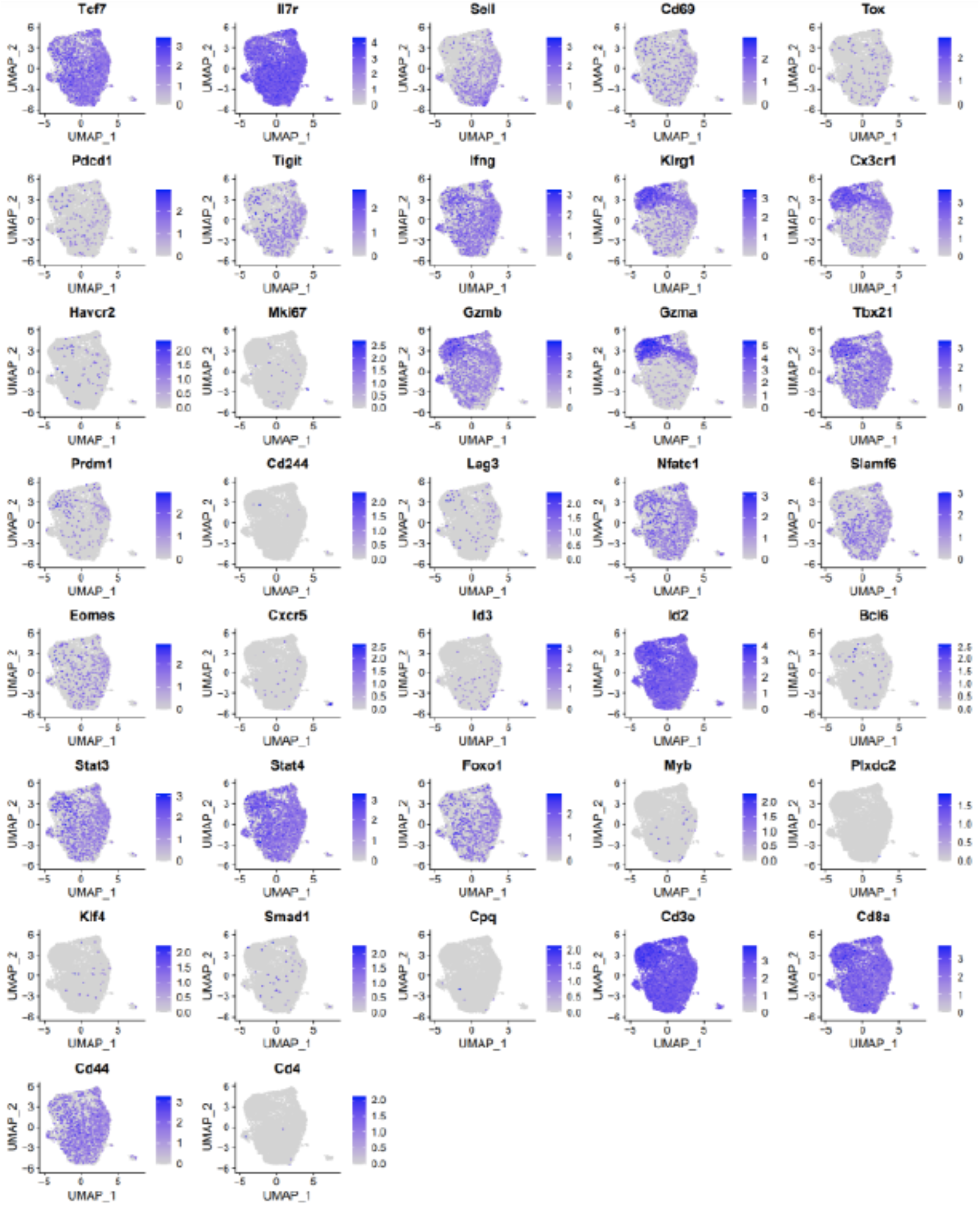
Normalized gene expression for select genes of interest. All cells from two mice infected with acute LCMV were integrated into a single uniform manifold approximation projection (UMAP).

**Figure S13.**
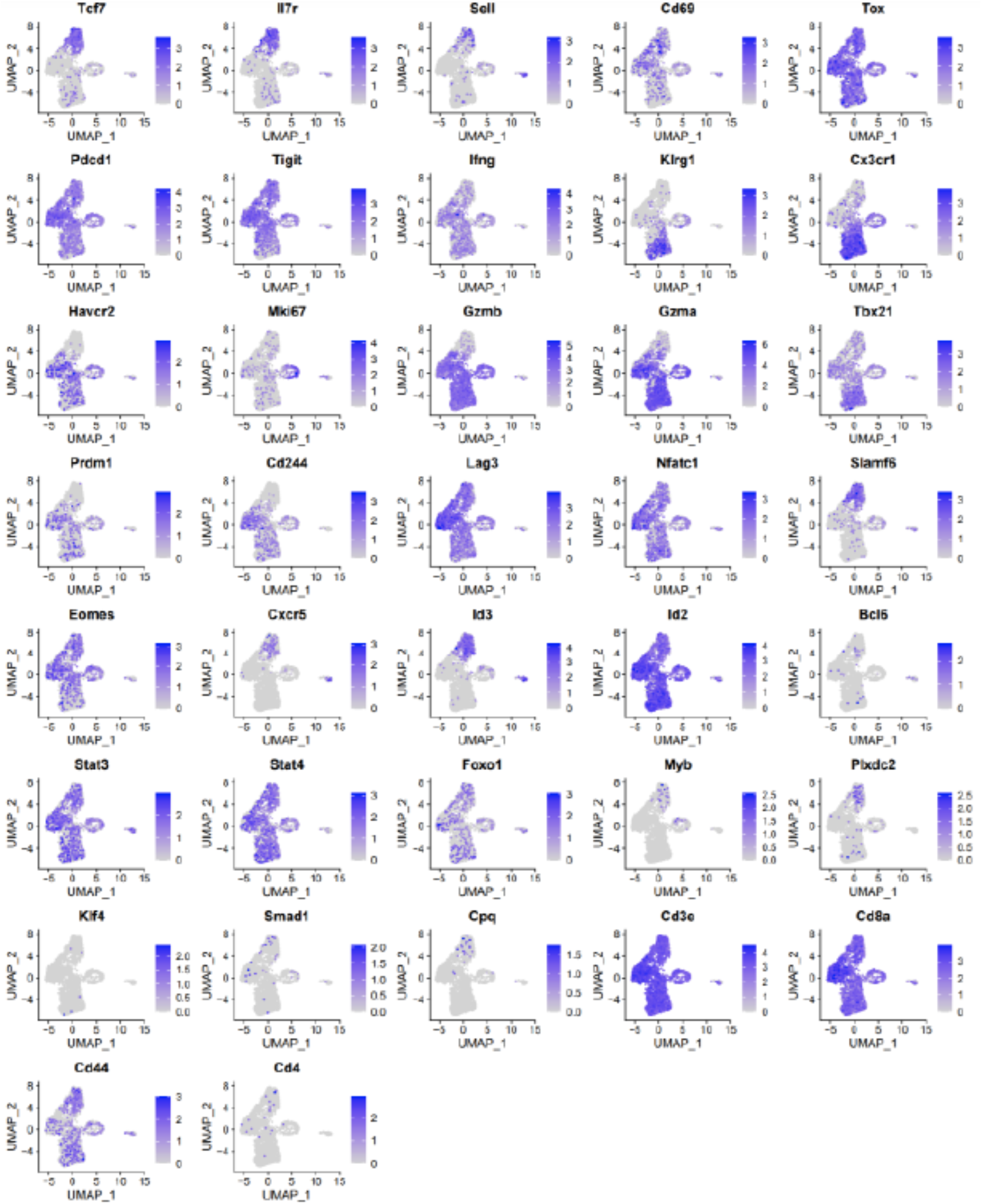
Normalized gene expression for select genes of interest. All cells from two mice infected with chronic LCMV were integrated into a single uniform manifold approximation projection (UMAP).

**Figure S14.**
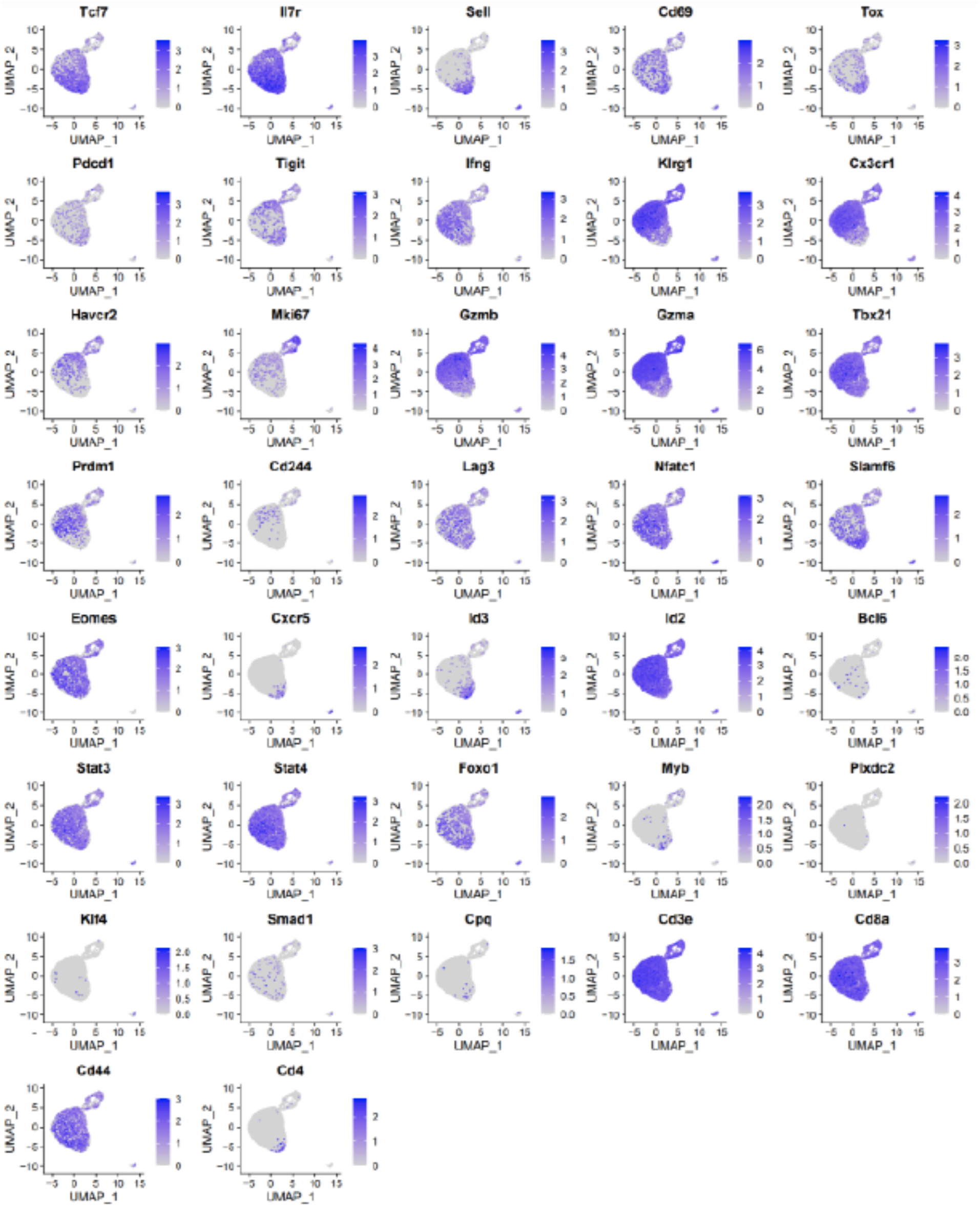
Normalized gene expression for select genes of interest. All cells from two mice infected with MCMV-*ie2*-gp33 were integrated into a single uniform manifold approximation projection (UMAP).

**Figure S15.**
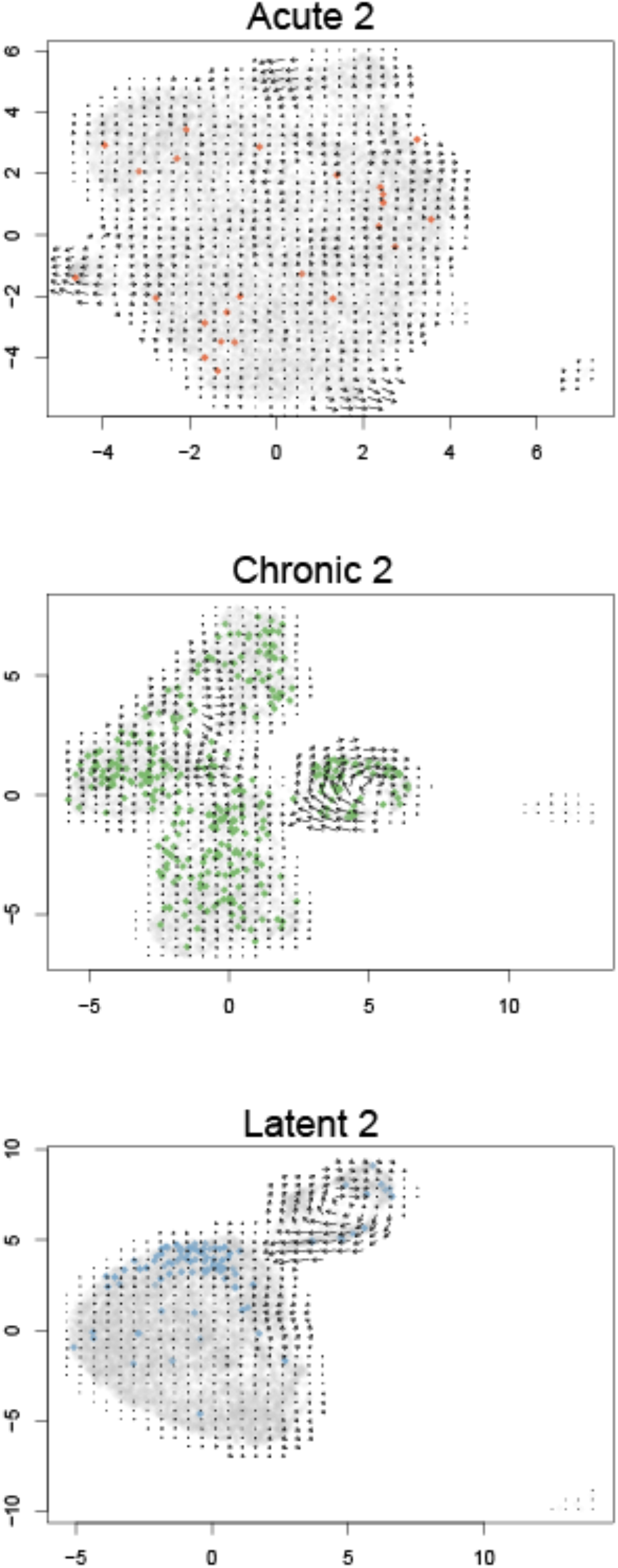
Pseudotime vector fields for each of the infection conditions. Colored points correspond to the most expanded clone found in a single mouse per infection type (Acute 2, Chronic 2, Latent 2). Clone was defined by identical CDRb3-CDRa3 nucleotide sequence.

**Figure S16.**
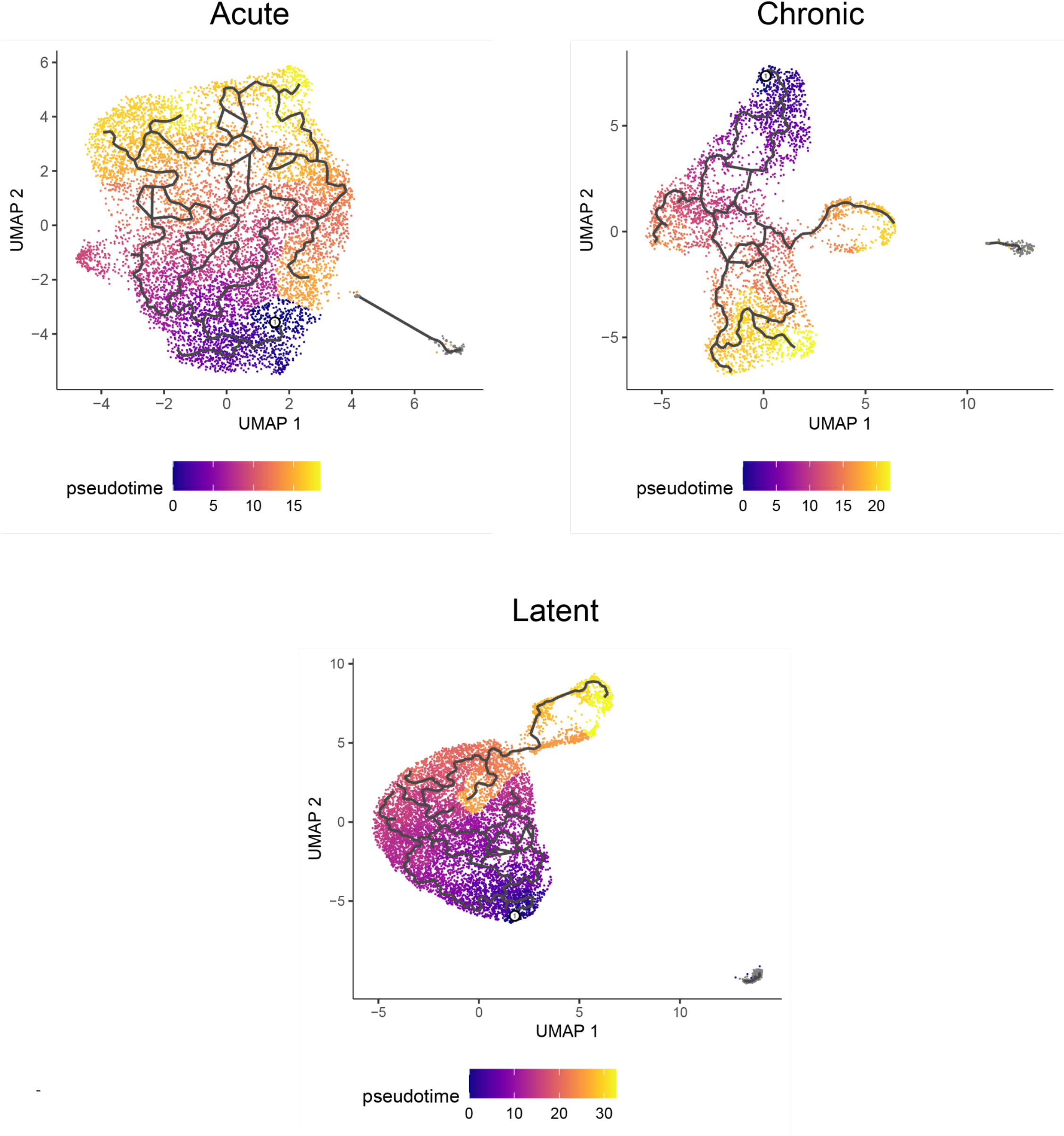
Uniform manifold approximation projections (UMAP) for cells isolated from mice infected with acute LCMV (acute), chronic LCMV (chronic), or MCMV-*ie2*-gp33 (latent). Each point indicates a cell and intensity corresponds to Monocle-calculated pseudotime. Solid lines indicate pseudotime trajectories.

**Figure S17.**
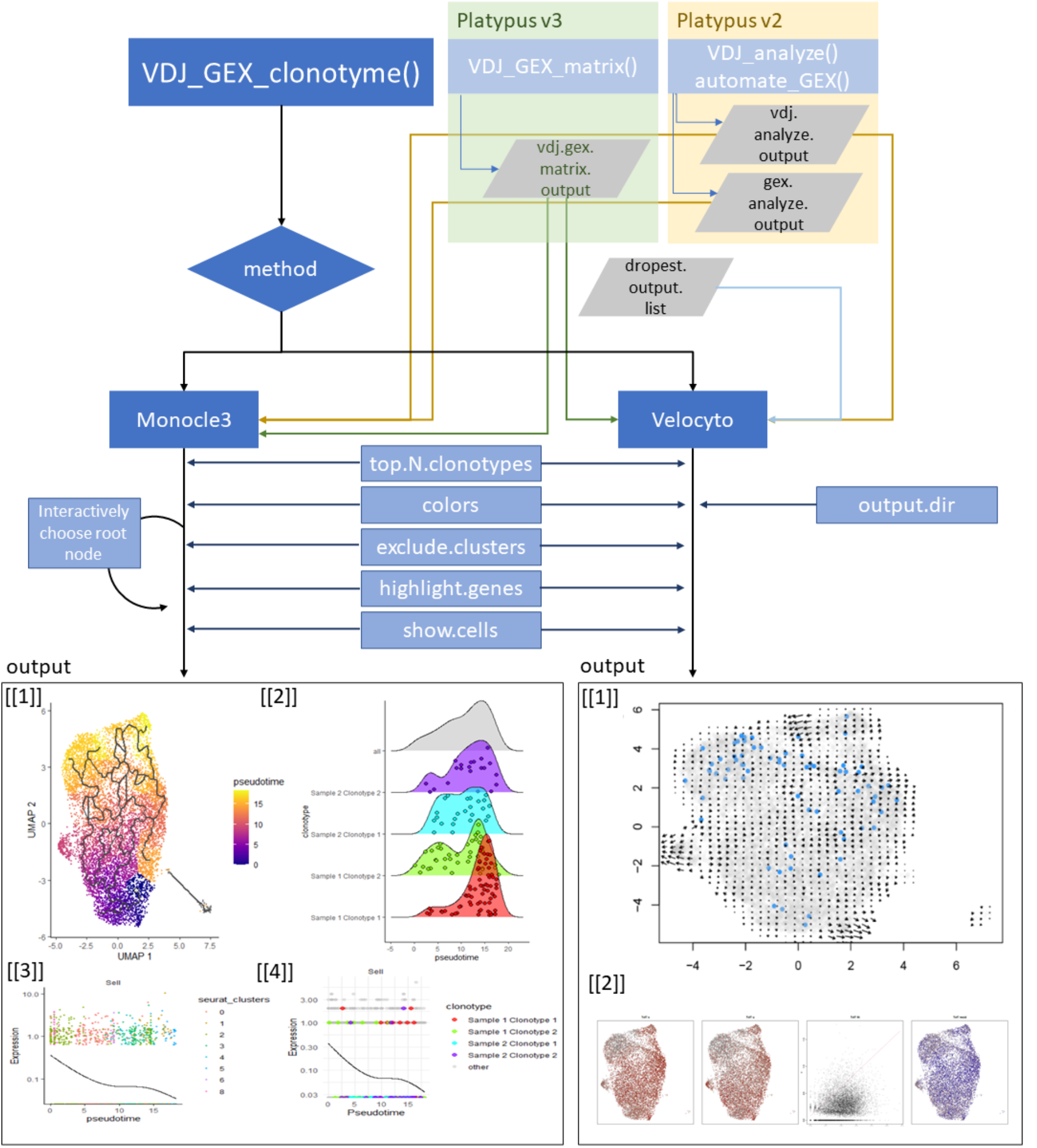
Clonotyme workflow. Clonotype information can be integrated with Monocle3 or Velocyto frameworks to investigate either global pseudotime properties or individual genes.

**Table S1:**
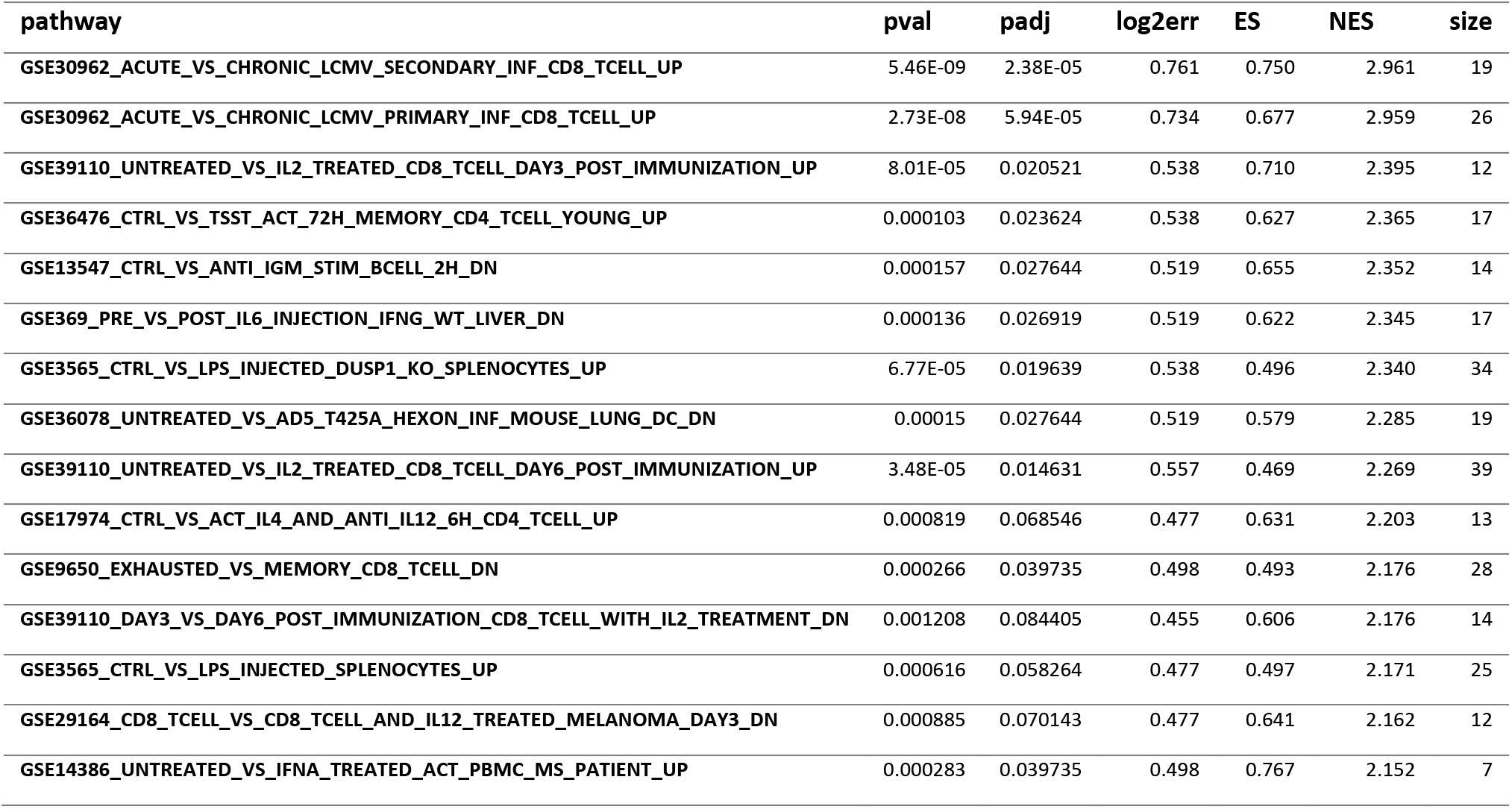
Top 15 gene sets based on the C7 immunological signatures from the Broad institute. The upregulated genes from MCMV-ie2-gp33 versus chronic LCMV infection were supplied as input.

